# The genome of Mekong tiger perch *(Datnioides undecimradiatus)* provides insights into the phylogenic position of Lobotiformes and biological conservation

**DOI:** 10.1101/787077

**Authors:** Shuai Sun, Yue Wang, Xiao Du, Lei Li, Xiaoning Hong, Xiaoyun Huang, He Zhang, Mengqi Zhang, Guangyi Fan, Xin Liu, Shanshan Liu

## Abstract

Mekong tiger perch (*Datnioides undecimradiatus*) is one ornamental fish and a vulnerable species, which belongs to order Lobotiformes. Here, we report a ∼595 Mb *D. undecimradiatus* genome, which is the first whole genome sequence in the order *Lobotiformes*. Based on this genome, the phylogenetic tree analysis suggested that *Lobotiformes* and *Sciaenidae* are closer than Tetraodontiformes, resolving a long-time dispute. We depicted the pigment synthesis pathway in Mekong tiger perch and result confirmed that this pathway had evolved from the shared whole genome duplication. We also estimated the demographic history of Mekong tiger perch, showing the effective population size suffered a continuous reduction possibly related to the contraction of immune-related genes. Our study provided a reference genome resource for the *Lobotiformes*, as well as insights into the phylogeny of Eupercaria and biological conservation.

## Instruction

Mekong tiger perch *(Datnioides undecimradiatus*) [1] is one tropical freshwater fish, belonging to the order Lobotiformes under series Eupercaria. It is native to Mekong river and usually found in the main waterway and large tributaries of the Mekong river basins, feeding on small fishes and shrimps [2]. It is also one ornamental fish, which is kept for its vertical yellow and black stripes running its body.

Eupercaria is by far the largest series of percomorphs with more than 6,600 species arranged in 161 families and at least 16 orders. The phylogenetic relationship of the order Lobotiformes, Tetraodontiformes, and the family Sciaenidae is in conflict. Mirande reported Sciaedidae as the sister clade of Tetraodontiformes, and then followed by Lobotiformes based on 44 DNA makers from uncompleted nuclear and mitochondrial sequences combined with morphological characters [3]. Compared to it, Betancur-R et al. reported Lobotiformes was more closely related to Tetraodontiformes than Sciaedidae using molecular and genomic data, which was also not complete or whole-genome sequenced for most of species. [4], agreed with this phylogeny by investigating. However, more recently Lobotiformes was reported to be more closely related to Sciaenidae than Tetraodontiformes based on complete mitochondrial genome [5]using and transcriptomic data [6]. Furthermore, fourteen families of Eupercaria included in order-level *incertae sedis*, which are called “new bush at the top”, were not arranged to explicit orders and interrelationships among them was a long-term issue [7]. Therefore, the whole-genome containing comprehensive evolutionary information is called for resolving the long-time dispute on the phylogenetic relationships of the huge number of species in Eupercaria, especially for the problem of “new bush at the top”.

In addition to its evolutionary importance, Mekong tiger perch has a body color pattern with vertical yellow and black stripes. Body color diversity in animals has important functions in numerous biological processes and social behaviors, such as sexual selection, kin recognition and changing coloration for camouflage [8]. Recent studies proposed that teleost genomes might contain more copies of genes involved in pigment cell development than tetrapod genomes after an ancient fish-specific genome duplication (FSGD), which might contribute to the evolution and diversification of the pigmentation gene repertoire in teleost fish [9]. With more genome sequences, especially for fish with unique body color schemes such as Mekong tiger perch, we can further apply comparative genomics to illustrate the genetic mechanisms of body color development.

Mekong tiger perch is currently assigned as ‘Vulnerable’ due to the rapidly declined population size by the IUCN [10], and is considered as endangered (EN) by Thailand Red data [2]. The external factors, such as the construction of hydraulic engineering infrastructures, urban pollution, and the aquarium trade, are thought to be exerting a negative effect on wild populations. Meanwhile, internal genetic factors such as resistance to biological and abiotic stress may be related to their survivals. Due to its limited distribution and commercial values, rare genetic research has been focused on Mekong tiger perch. With the rapid development of genomics, each fish deserves the right to own its genome assembly representing its unique genetic resource, which will help to better investigate its unique characters and biological conservations.

Here, we sequenced Mekong tiger perch and assembled a reference genome, which was the first genome of the order Lobotiformes. We constructed a phylogenetic tree in Eupercaria based on the whole genome sequences, to elucidate the relationships among family Sciaenidae, order Lobotiformes and order Tetraodontiformes, providing insights into the phylogenic position of Lobotiformes. Utilizing the assembled genome, we identified genes involved in the cell development regulation and pigment synthesis in Mekong tiger perch. We confirmed population decline by the analysis of demographic history and found the contraction of immune-related genes might be a contributing factor for Mekong tiger perch’s vulnerability. The genome assembly of Mekong tiger perch provided a valuable genome resource for future fish studies in Lobotiformes, and also contributes to the understanding of body color development as well as demographic history and conservation.

## Results

### Genome assembly, annotation, and genomic features

We sampled muscle tissue from a Mekong tiger perch captured in Mekong river (Supplementary Fig. 1), and applied single tube long fragment read (stLFR) [11] technology for whole genome sequencing, generating stLFR co-barcode reads 122.4 Gb raw data. After filtering low-quality and duplicated reads, we obtioned75.3 Gb clean data for genome assembly using supernova [12], and gaps were closed using GapCloser [13]. We obtained a final genome assembly spanning 595 Mb, accounting for 95.5% of the estimated genome size (623 Mb, Supplementary Fig. 2). The assembly achieved a high level of contiguity, with a total of 4,959 scaffolds and scaffold N50 of 9.73Mb. The longest 71 scaffolds (longer than 1.41 Mb) accounted for 90% of the total genome, and the longest scaffold reached up to 39.31 Mb (Fig. 1, Table 1 **and** Supplementary table 1). Total repeat content accounted for 11.97% (Table 1 and Supplementary table 2) of the genome, and 29,150 protein-coding genes were predicted via *ab initio* and homology-based methods (Table 1). The average length of coding sequences (CDS) was 1,510 bp with an average of 9 exons per gene, which were similar to that of other related species (Supplementary table 3 **and** Supplementary Fig. 3). The ncRNAs including miRNA, tRNA, rRNA, and snRNA were also annotated with a total length of 179 kb (Supplementary table 4). We used BUSCO metazoan database (v9) to evaluate the completeness of gene sets and observed a completeness of 95.34%. Other databases including Actinopterygii (v9) and vertebrata (v9) estimated the completeness to be 94% and 91%, separately (Supplementary Fig. 4). Furthermore, the mitochondrial genome was assembled a total length of 16,606 bp, containing 18 coding genes, 2 rRNA, and 17 tRNA (Supplementary table 5).

**Table 1.** Assembly and annotation of the Mekong tiger perch genome

| Genome assembly |  | Repeat content |  | Genes finding and annotation |  |
| --- | --- | --- | --- | --- | --- |
| Total Length (bp) | 594,964,832 | DNA (%) | 6.11 | gene number | 29,150 |
| Number of scaffolds | 4442 | LINE (%) | 2.75 | average gene length (bp) | 13,759 |
| longest scaffold | 39.31 Mb | SINE (%) | 0.19 | average mRNA length (bp) | 1510 |
| N content (%) | 2.17 | LTR (%) | 2.37 | average exon number per gene | 9.04 |
| GC content | 42.74 | Other (%) | 0 | average exon length (bp) | 167.03 |
| N50 / NG50 | 9.73Mb / 18 | Unknown (%) | 3.35 | average intron length (bp) | 1522.83 |
| N90 / NG90 | 1.41Mb / 71 | Total (%) | 11.97 | annotated genes | 19,837 |

CpG islands (CGIs) are an important group of CpG dinucleotides in the guanine- and cytosine rich regions as they harbor functionally relevant epigenetic loci for whole genome studies. 32,148 CpG islands (CGIs) were identified with a total length up to 18.8 Mb. The CpG density has the most prominent correlations with three other genomic features. It correlated positively with CG content density and gene density, and correlated negatively with repeat density (Fig. 1b **and** Supplementary table 6), showing a similar pattern observed in other published fishes or mammal [14–16].

### Phylogenetic tree of Eupercaria uncovers the phylogenetic position of Lobotiformes

To clarify the evolutionary relationships of major orders in Eupercaria clade, 9 sequenced species from 7 different orders were used in the comparative genomics analysis (Supplementary table 7). We clustered the gene families based on protein sequences similarity and obtained a total of 13,785 gene families, 1,428 of which were single-copy gene families (Supplementary Fig. 5 **and** Supplementary table 8). The nucleotide sequences on the four-fold degenerate (4d) site of those single-copy gene families were used to construct the maximum likelihood (ML) tree. The phylogeny of the seven orders was found consistent with the previous study [5, 6]. Order Perciformes was identified as the early branch to other orders in Eupercaria, and the divergent time was estimated 101.9 million years ago (mya). Our phylogenetic tree (Fig. 2a) showed Lobotiformes was more related to Sciaenidae than to Tetraodontiformes, which supported the results of some previous studies [5, 6]. Furthermore, the divergence time between Lobotiformes and Sciaenidae was inferred to be 82.9 mya (Fig 2a).

**Fig 1.**
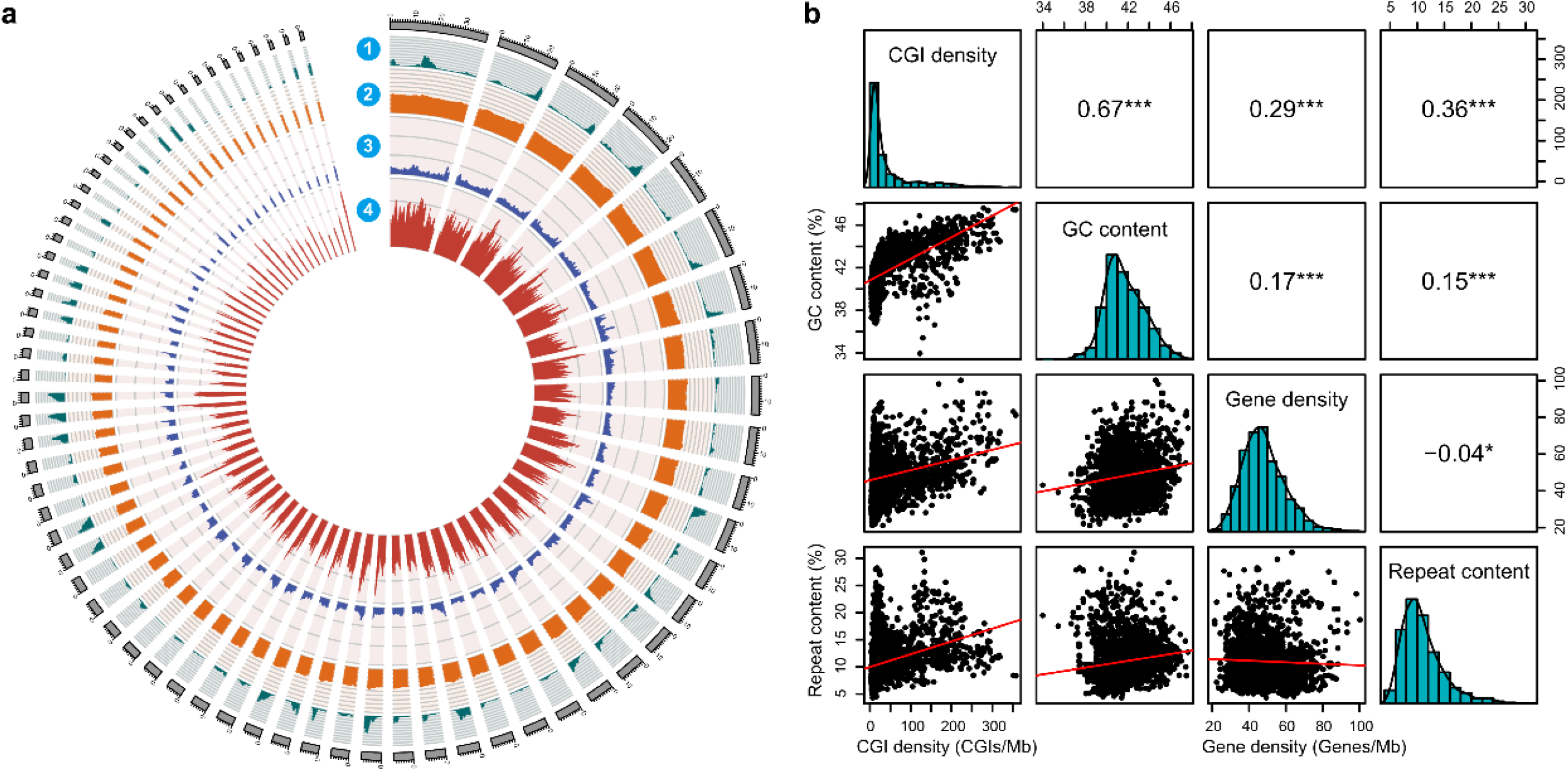
The genome features of *D. undecimradiatus.* **(a)** A Circos plot representing four features using sliding overlapping windows of 1Mb length with 200kb step through the 71 scaffolds (scales in Mb), which accounts for 90% of genome. (1) CGI content, measured by CGIs number per million base pairs (megabase, Mb). The range of the axis is 0 to 500. (2) GC content, measured by the proportion of GC in unambiguous bases of 1 Mb window size. The range of the axis is 0 to 100. (3) Repeat content, measured by the proportion of repeat regions of 1 Mb window size. The range of the axis is 0 to 100. (4) Gene density, measured by genes number per million base pairs. **(b)** Correlation matrix plot with significance levels between four genome features. The lower triangular matrix is composed by the bivariate scatter plots with a fitted linear model. The diagonal shows the distribution by histogram with density curve. The upper triangular matrix shows the Pearson correlation plus significance level (as stars). Different significance levels are highlighted with asterisks: p-values 0.001 (***), 0.01 (**), 0.05 (*). This plot was generated with the “psych” package in R (v3.5.0).

**Fig. 2.**
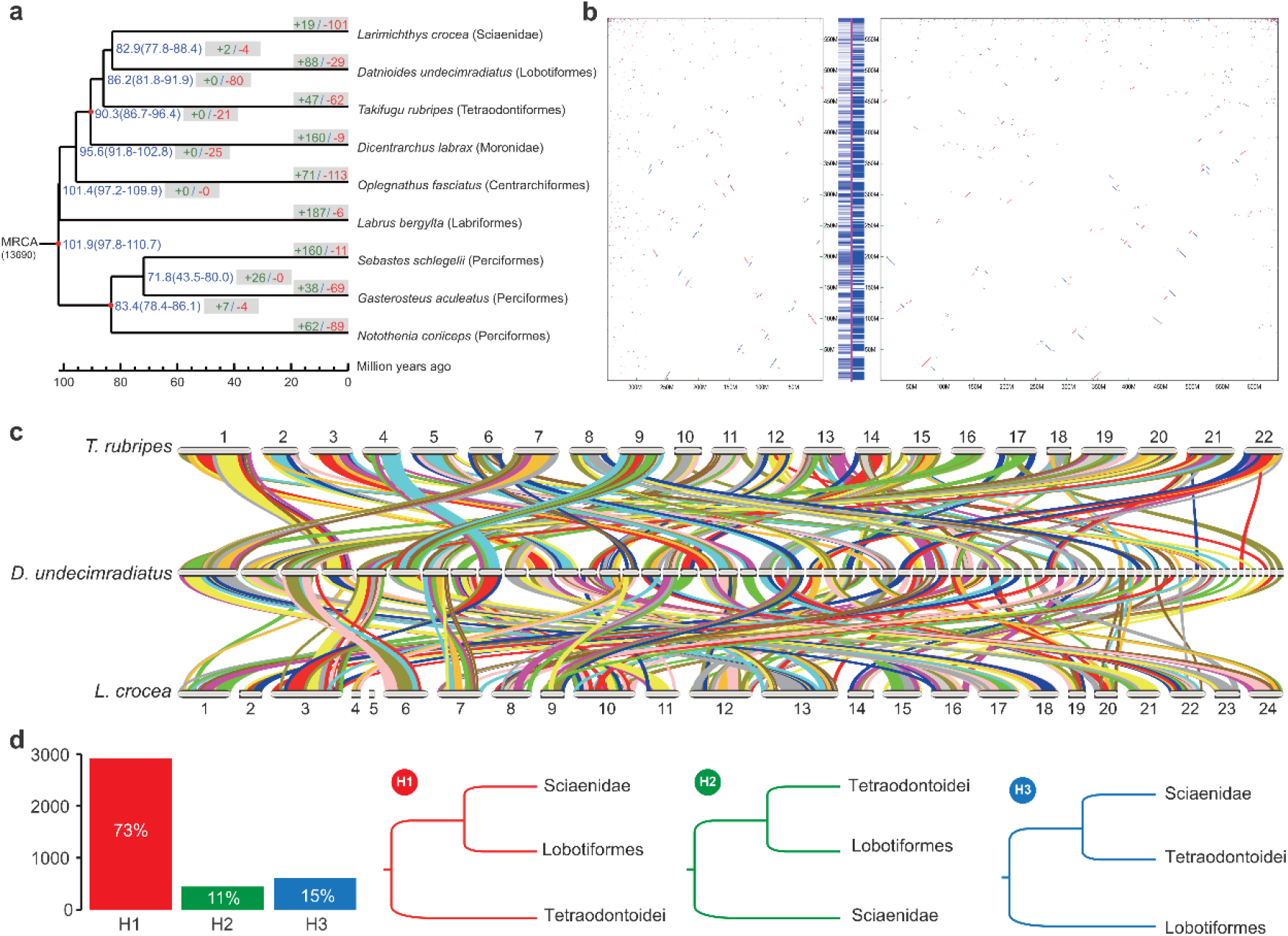
The phylogenetic and genome analysis for *D. undecimradiatus* and other related species. **(a)** Time-calibrated maximum likelihood phylogenetic tree of nine species from Eupercaria. Red Nodes represents the calibration time points obtained from TreeTime. The estimated divergent time (mean and 95% highest-probability) are showed on the right of the nodes. Behind the divergent time, the green positive number and the red negative number in grey boxes stand for the number of gene families significantly expanded and contracted, respectively. **(b)** Synteny of *D. undecimradiatus* between *L. crocea and T. rubripes* at whole-genome nucleotide level. The y-axis represents the *D. undecimradiatus* genome, and the left x-axis refers to *T. rubripes* and right x-axis refers to *L. crocea*. The fringe plot on the left of y-axis represents the synteny regions between *D. undecimradiatus* and *T. rubripes* on *D. undecimradiatus* genome. The fringe plot on the right of y-axis represents the synteny regions between *D. undecimradiatus* and *L. crocea* on *D. undecimradiatus* genome. **(c)** synteny of *D. undecimradiatus* between *L. crocea and T. rubripes* at gene level and different colors represent different synteny blocks. **(d)** The number of gene genealogy that supports for three hypotheses concerning the Lobotiformes phylogeny.

The conserved genomic synteny reflects the arrangements on evolutionary processes was used to further demonstrate the ambiguous phylogeny among Lobotiformes and closely related orders. The syntenic analysis both at whole-genome nucleotide-level and gene-level were performed by aligning *L. crocea* and *T. rubripes* to our assembled *D. undecimradiatus*, separately. At whole-genome nucleotide-level after filtering alignment length < 1kb, 41.76 % of *D. undecimradiatus* genome sequences were covered by *L. crocea* genome with an average of 2.29 kb per block. In comparison, only 10.48% of *D. undecimradiatus* genome sequences were covered by *T. rubripes* genome with an average of 1.65 kb per block (Fig. 2b **and** Supplementary table 9). Similarly, at the whole genes level after keeping block length > 3 genes, 89.19% of *D. undecimradiatus* genes showed synteny with *L. crocea* with an average of 50.28 genes per block, and 77.21% of *D. undecimradiatus* genes had synteny with *T. rubripes* with an average of 41.28 genes per block (Fig. 2c **and** Supplementary table 10). Despite the difference in genome size, both *L. crocea* and *T. rubripes* genomes were assembled to comparable chromosome level and the BUSCO assessments showed no significant differences in the completeness of genome and gene set between *L. crocea* and *T. rubripes* (Supplementary table 11). In addition, the distribution of the length of syntenic blocks at both whole-genome nucleotide-level and gene-level showed significant difference by t-test statistics (nucleotide-level, *p*-value<0.0001; gene-level, *p*-value<0.05) (Supplementary Fig. 6). Therefore, the results of synteny suggested that the Lobotiformes had better evolutionary conservation and closer relationship with Sciaenidae than with Tetraodontiformes, providing strong evidence for the constructed phylogenic tree.

To further demonstrate the phylogeny among Lobotiformes, Sciaenidae, and Tetraodontiformes, only the homologous genes of 1:1:1 on the syntenic blocks were inferred as reliable orthologous genes to construct the genes genealogy, with the human CDS sequences as outgroup in the rooted tree. 73% of the 3,974 orthologous gene sets supported that *D. undecimradiatus* was more closely related to *L. crocea*, supporting Lobotiformes as sister group to Sciaenidae, instead of the other two hypotheses (Fig. 2d).

### The genes involved in pigment development and two copies of three rate-limiting genes of pigment synthesis as the result of FSGD similar to other teleosts

In consideration of special skin color pattern, among established pigmentation database containing 198 genes [17], 172 genes were found on our genome, occupying 92% of the database and possibly genetic basis for the phenotypic characteristics of vertical yellow and black stripes running its body (Supplementary table 12). For two major pigment synthesis pathways in *D. undecimradiatus*, three main rate-limiting genes of Tyrosinase family (*TYR*, *DCT*, *TYRP1*) in the melanin synthesis pathway have two copies respectively (Fig.3a) and one main rate-limiting gene SPR in pteridine synthesis pathway also have two copies (Fig. 3b). This suggests *D. undecimradiatus* retained the pigment related genes from a fish-specific whole-genome duplication (FSGD), which was also observed in many other teleosts. It should be noted that the phylogeny of those genes also served as another evidence to the evolution relationship among the three close relative orders.

**Fig. 3.**
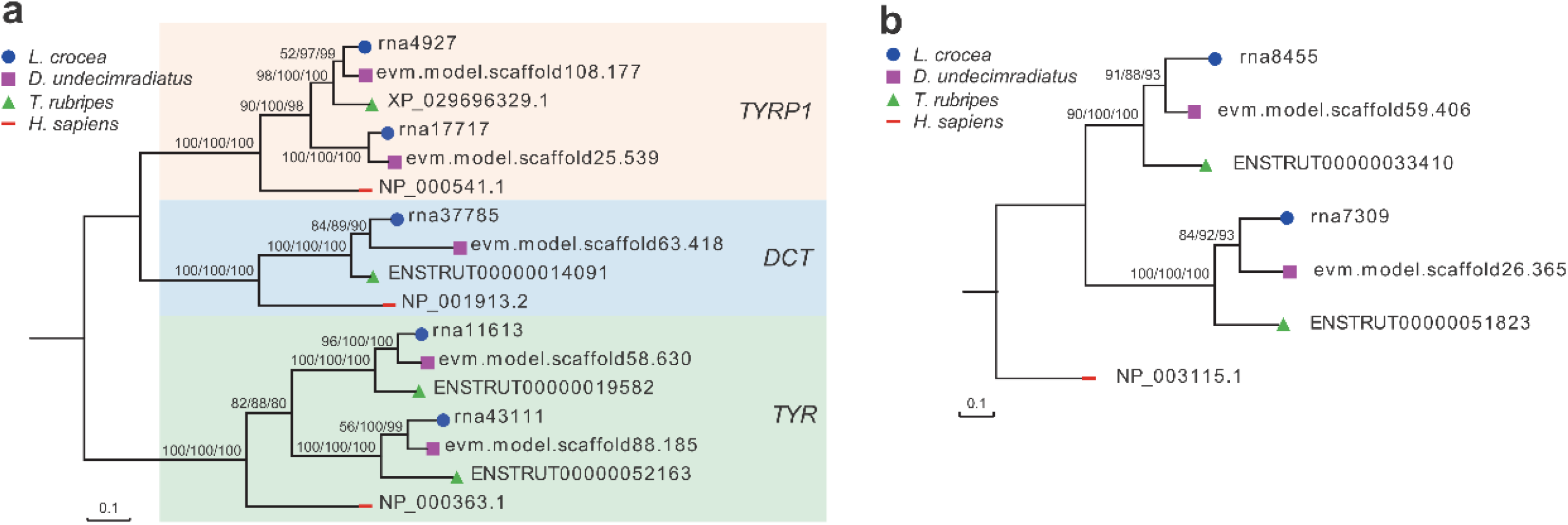
The phylogeny of main rate-limiting genes involved in the pigment synthesis. **(a)** The concordant phylogeny of TYR gene family (*TYRP1*, *DCT* and *TYR*) using maximum likelihood, neighbor-join and minimal evolutionary methods, Corresponding bootstrap values were showed on branch labels. **(b)** The concordant phylogeny of *SPR* gene using maximum likelihood, neighbor-join and minimal evolutionary methods. Corresponding bootstrap values were showed on branch labels.

### Decreasing population size related to the contraction of immune-related gene families provides clues to biological conservation

Pairwise sequentially Markovian coalescent (PSMC) was used to infer the demographic history of Mekong tiger perch. The effective population size continuously reduced since LGM (last glacial maximum), and there were no signs of recovery to date, which was consistent with its vulnerable state [2, 10](Fig. 4a).

**Fig. 4.**
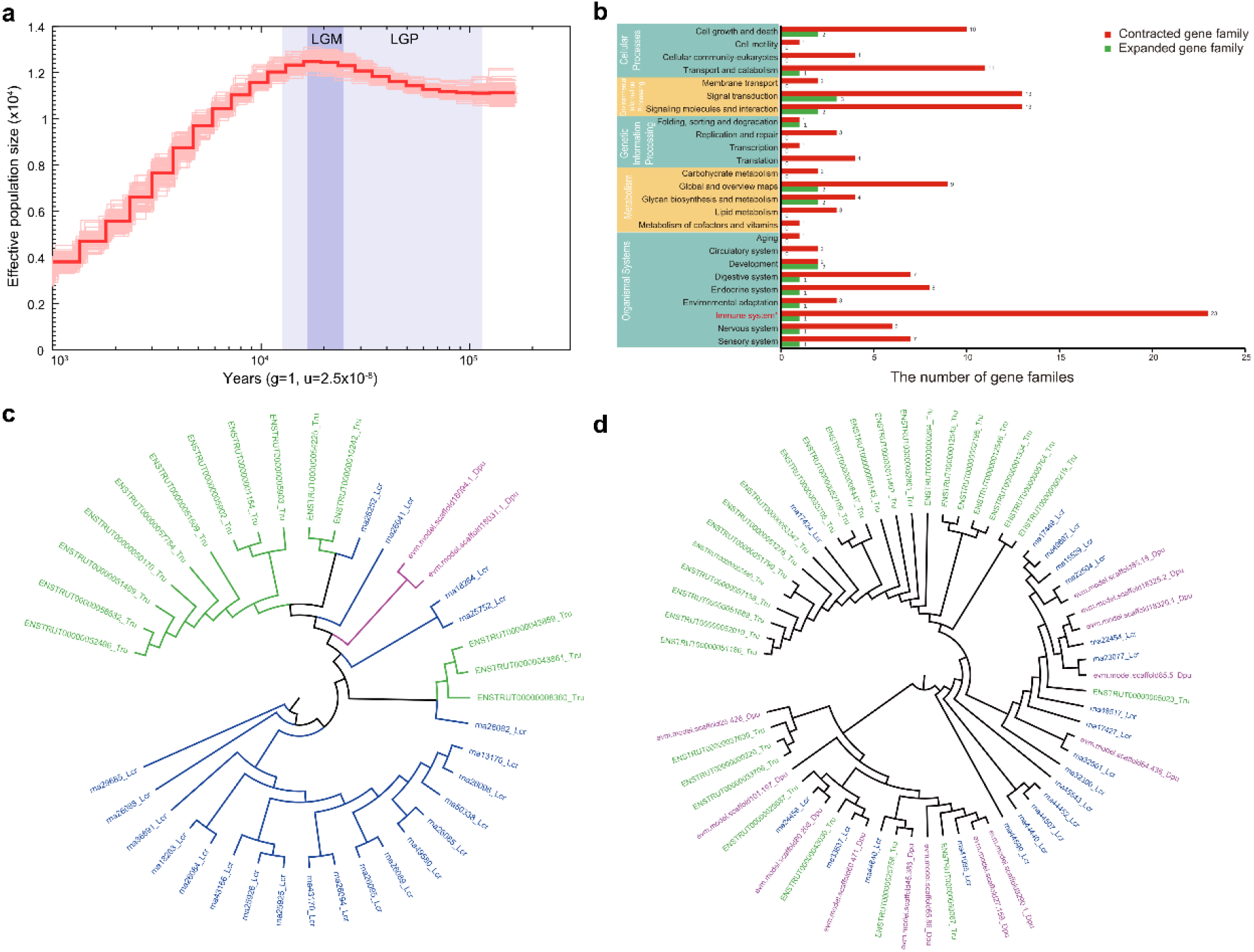
The demographic history of *D. undecimradiatus* and genetic basis possibly associated with vulnerability. **(a)** The population history of *D. undecimradiatus* inferred using PSMC. LGM (last glacial maximum, ∼26.5-19 kya) and LGP (last glacial period, ∼102-1.05 kya) are shaded in gray. **(b)** The number of significantly contracted and expanded gene families (*P*-value < 0.05) involved in different KEGG pathway (at level 1 and level 2). The number at the right end of the bar indicates the number of gene families. **(c)** Phylogenetic tree of *MHC1* gene family. Green color refers to *T. rubripes*, and blue and red refers to *L. crocea* and *D. undecimradiatu* separately. **(d)** Phylogenetic tree of *NLRP12* gene family. Green color represents *T. rubripes*, and blue and red represents *L. crocea* and *D. undecimradiatu* separately.

The change of genes copy number plays a role in the species adaptation [18] To investigate the genetic basis potentially related to fish survival, we identified the expanded and contracted gene families in Mekong tiger perch, and 19 and 101 significantly expanded and contracted gene families were found (*p*-value < 0.05), separately (Fig. 4b). 62 contracted and 18 expanded gene families were annotated to KEGG ortholog functions. The top enriched KEGG pathway was the immune-related pathway with fifteen contracted gene families (Fig. 4c **and** Supplementary table 13-14), Furthermore, the most gene families’ KO were annotated to the genes MHC1, NLRP12, ANK (ankyrin), IGH CLDN, and PLAUR (Supplementary Table 15), which may play a role in the adaptive immunity and survival. For example, MCH1, which was responsible for presenting peptides on the cell surface for recognition by T cells [19], was significantly contracted in *D. undecimradiatus* with only 2 copies, compared to 22 copies in closely related *L. crocea* and 14 copies in *T. rubripes* (Fig. 4c). NLRP12, which play a role in regulating inflammation and immunity [20], there were 13 copies in *D. undecimradiatus*, compared to 20 copies in *L. crocea and 29 copies in T. rubripes.* The contraction of immune-related genes may affect the capacity of Mekong tiger perch to adapt to environmental changes or stress, implying of human-mediated species and environmental conservation is meaningful.

## Discuss

The phylogeny of Eupercaria plays a fundamental role in species classification and uncovering the species evolutionary history at the Cretaceous–Palaeogene boundary[21]. However, although species in series Eupercaria account for more than twenty percent of the bony fish, the Eupercaria phylogeny is ambiguous or conflicted, especially for the “new bush at the top”. Meanwhile, the resolution of the phylogeny is currently limited to the order level, and few studies could go down to the class level or species level. Relying on limited morphological characters and molecular sequences, it is difficult to draw convincing conclusion[3–6]s. In contrast, whole genome sequencing provides sufficient information to perform the phylogenetic analysis of species. In our study, we clarified the relationship of order Lobotiformes to its related orders, Sciaenidae and Tetraodontiformes. With the rapid development of sequencing technology, large-scale fish genome sequencing projects can be achieved, such as fish 10k project (http://icg-ocean.genomics.cn/index.php/fish10kintroduction/). Those projects will greatly promote studies on the fishes’ classification and evolution.

Skin color is a biologically important trait, which is a fascinating research topic and has great implications on biological adaption, commercial value, and skin health[22, 23]. In our study, most genes involved in pigmentation development, regulation and synthesis can be found in our assembled genome. However, research of underlying mechanisms is difficult to penetrate with limited genome resource. More fish genome data and molecular experiments will facilitate the analysis of skin color regulation mechanisms.

Biological conservation is an important research content of the relationship between human and nature [23], and different species vary in their adaptive capacities. In our study, immune-related genes of Mekong tiger perch were significantly contracted relative to closely related species, indicating that its environmental adaptability possibly responsible for its vulnerability. Therefore, it is necessary for humans to take various measures to protect it, such as improving the living environment, reducing fishing, artificial breeding, etc., and thus help maintain species diversity.

## Materials and Method

### DNA extraction and stLFR library construction, and sequencing

The long genomic DNA molecules were extracted from the muscle of Mekong tiger perch. The stLFR library was constructed following the standard protocol using MGIEasy stLFR library preparation kit (PN:1000005622) [11]. In details, the transposons with hybridization sequences were inserted in the long DNA molecules approximately every 200-1000 base pairs. The transposon integrated DNAs was then mixed with beads that each contained an adapter sequence. A unique barcode was shared by all adapters on each bead with a PCR primer site and a capture sequence that is complementary to the sequences on the integrated transposons. When the genomic DNA was captured to the beads, the transposons were ligated to the barcode adapters. After a few additional library-processing steps, the co-barcoded sub-fragments were sequenced on the BGISEQ-500 sequencer.

### Reads filtering, genome size estimation and genome assembly

We generated a total of 1,223,801,322 million raw pair-end co-barcoding reads of 122.4 Gb. To obtain high-quality genome, SOAPnuke (v2.2) [24]was performed to filter low-quality reads (>40% low-quality bases, Q<7), PCR duplications, or adapter contaminations. After that, 753,357,182 clean pair-end reads remained. Based on the 17-mer analysis, the Mekong tiger perch genome size was estimated to be 623 Mb. Supernova assembler v 2.0.1(10X Genomics, Pleasanton, CA) were used to construct contig and scaffold, followed by gaps closing using GapCloser (v1.2) [13].

### Genes structure and function annotation

We used both *de novo* approaches and homology-based approaches to predict repeat elements in *D. undecimradiatus* genome. Firstly, we aligned our genome against the Repbase database [25] at both protein and DNA levels by using RepeatMasker (v4.0.5) and RepeatProteinMasker (v4.0.5) [26] to identify transcriptional elements (TEs). Secondly, we used RepeatModeler (v1.0.8)[27] and LTR-FINDER (v1.0.6)[28] to implement *de novo* repeat annotation. Next, we used RepeatMasker to complete repeat elements identification and classification. Lastly, we combined and classified the above results. We masked the repeats in *D. undecimradiatus* genome for the subsequent gene finding.

For homology-based annotation, we downloaded protein sequences of *Dicentrarchus labrax*, *Labrus bergylta*, *Larimichthys crocea*, and *Gasterosteus aculeatus* from NCBI (https://www.ncbi.nlm.nih.gov/) and Ensembl (http://ensemblgenomes.org/). We aligned these sequence to *D. undecimradiatus* genome using BLAST[29] with an E-value cutoff of 1e^−5^ and the matched length coverage >30% to identify homologous genes. Based on the aligned result, we used GeneWise (v2.4.1) [30] to predict gene models. Furthermore, we used AUGUSTUS (v3.1) [31]and GENSCAN(v2009) [32] for *de novo* annotation with default parameters and zebrafish data setting as a training set of AUGUSTUS. Lastly, we integrated all above gene models by EVM [33]. We used BUSCO (v 3.0.2) [34] to assessment gene annotation integrity with three different databases, including vertebrata (v9), metazoan (v9), and vertebrata (v9).

### Gene functional annotation

In order to perform gene functional annotation, we aligned above gene sets against Kyoto Encyclopedia of Genes and Genome (KEGG v87.0) [35], and NR (v84)[36] databases by blastp (E-value≤1e-5) to identify genes with similar functions. For identifying gene motifs and domains and obtaining Gene ontology (GO) terms [37], we aligned our predicted genes against ProDom [38], Pfam [39], SMART [40], PANTHER [41], and PROSITE [42] using InterProScan [43].

### ncRNA annotation

Five types of ncRNA (Non-coding RNA), including tRNA, snRNA, miRNA, snRNA and, rRNA were predicted. We used tRNAscan-SE (v1.3.1) to predict tRNA in our genome with the default parameters. The genome was aligned against Rfam(v12.0) (Nawrocki E P et al., 2015) database and then we used infernal (v1.1.1) (Nawrocki E P & Eddy S R, 2013) to infer snRNA and miRNA based on mapping result. We aligned vertebrate rRNA database against *D.undecimradiatus* genome to predict rRNA.

### CpG islands identification

The CpG islands (CGIs), which are clusters of CpGs in CG-rich regions, were identified using CpGIScan with the parameters “--length 500 --gcc 55 --oe 0.65”[44].

### Comparative genome analysis

We download the annotation files of 8 species including *Dicentrarchus labrax*, *Gasterosteus aculeatus*, *Labrus bergylta*, *Labrus bergylta*, *Notothenia coriiceps*, *Oplegnathus fasciatus*, *Larimichthys crocea*, and *Takifugu rubripes* form NCBI or Ensembl database (Suppl. Table 5). The longest transcript was extracted for each gene. We filtered error sequences that didn’t have enough sequence length, with termination codon in the middle, and with sequence length not divisible by 3 to obtain raw gene sets for each species. TreeFam (v4.0) [45] was used to identify gene families.

We concatenated single-copy genes into a supergene for each species and identified fourfold degenerate sites within each supergene to construct a phylogenetic tree using RAxML (v8.2.12) [46] with GTRCATX nucleotide substitution model with parameters as “-f a -x 12345 -p 12345”. Then by using the split time between *Gasterosteus aculeatus* and *Larimichthys crocea*, *Dicentrarchus labrax* and *Larimichthys crocea*, and *Notothenia coriiceps* and *Gasterosteus aculeatus* from timetree (http://www.timetree.org/) [47] as the reference time points, we estimated the divergent time between each species by MCMCtree from the PAML package with default parameters [48].

### Gene family cluster and expansion and contraction analysis

Expansion and contraction of each gene family were identified by Café (v2.1) [49] based on divergence time tree. To obtain the potential functions of the gene families, the number of different KO terms was counted for each gene family. The functions of the gene families were assigned by the corresponding KO terms of more than half counts or the highest count. The KEGG pathways involved by KO terms were extracted for further functional analysis. For the phylogenetic tree of the gene family, the CDS sequences were fetched to construct ML tree using RAxML (v8.2.12) [46].

### Synteny analysis

The synteny analysis of *D. undecimradiatus* against *L. crocea* and *T. rubripes* was performed on both whole-genome nucleotide level and gene level. On nucleotide level, we used Lastz (v1.02.00) [50] to identify synteny blocks with parameters “T=2 C=2 H=2000 Y=3400 L=6000 K=2200”, and aligned blocks less than 1 kb were filtered.

On gene level, we used JCVI (v0.8.12) [51] to identify synteny genes in two combinations based on CDS. The JCVI carried out sequence alignment based on Lastal (v979) with parameters “-u 0 -P 48 -i3G -f BlastTab”. The JCVI filtered the blast result based on C-score (C-score(A,B) = score(A,B) / max(best score for A, best score for B)) with parameters “C-score >= 0.70 tandem_Nmax=10”. We also filtered out the blocks spanning less than 30 genes on the *D. undecimradiatus* genome.

### Population demographic history inference

The history of effective population size was reconstructed using PSMC (v0.6.5-r67) [52]. Diploid genome reference for the individual were constructed using SAMtools and BCFtools [53] with the parameters of “samtools mpileup -C30” and “vcfutils.pl vcf2fq -d 10 -D 100”. The demographic history was inferred using PSMC with “-N25 -t15 -r5 -p 4+25*2+4+6” parameters. The estimated generation time (g) and mutation rate per generation per site (μ) were set to 1 and 2.5e-8.

## Data available

The sequencing reads of Mekong tiger perch in this study have been deposited in NCBI Sequence Read Archive (SRA) under BioProject accession PRJNA574247. The datasets reported in this study are also available in the CNGB under accession number CNP0000691.

## Acknowledgements

This work was financially supported by the special funding of “Blue granary” scientific and technological innovation of China (2018YFD0900301-05) and Special Fund for Marine Economic Development of Fujian Province (ZHHY-2019-3). We also thanks for the technical support of stLFR library construction and sequencing from China National Genebank.

## Author Contributions

X. L., G. F. and H. Z. conceived the project. S. L. and G. F. supervised the study. M. Z. contributed to sample collections. S. S., Y. W., L. L., and X. H. performed bioinformatics analyses. S. S, Y.W., G. F., X. L., and X. D. wrote the manuscript with help from all co-authors.

## Competing Interests

The authors declare no competing interests.

## Supplementary Figures and Tables

**Supplementary figure 1.**
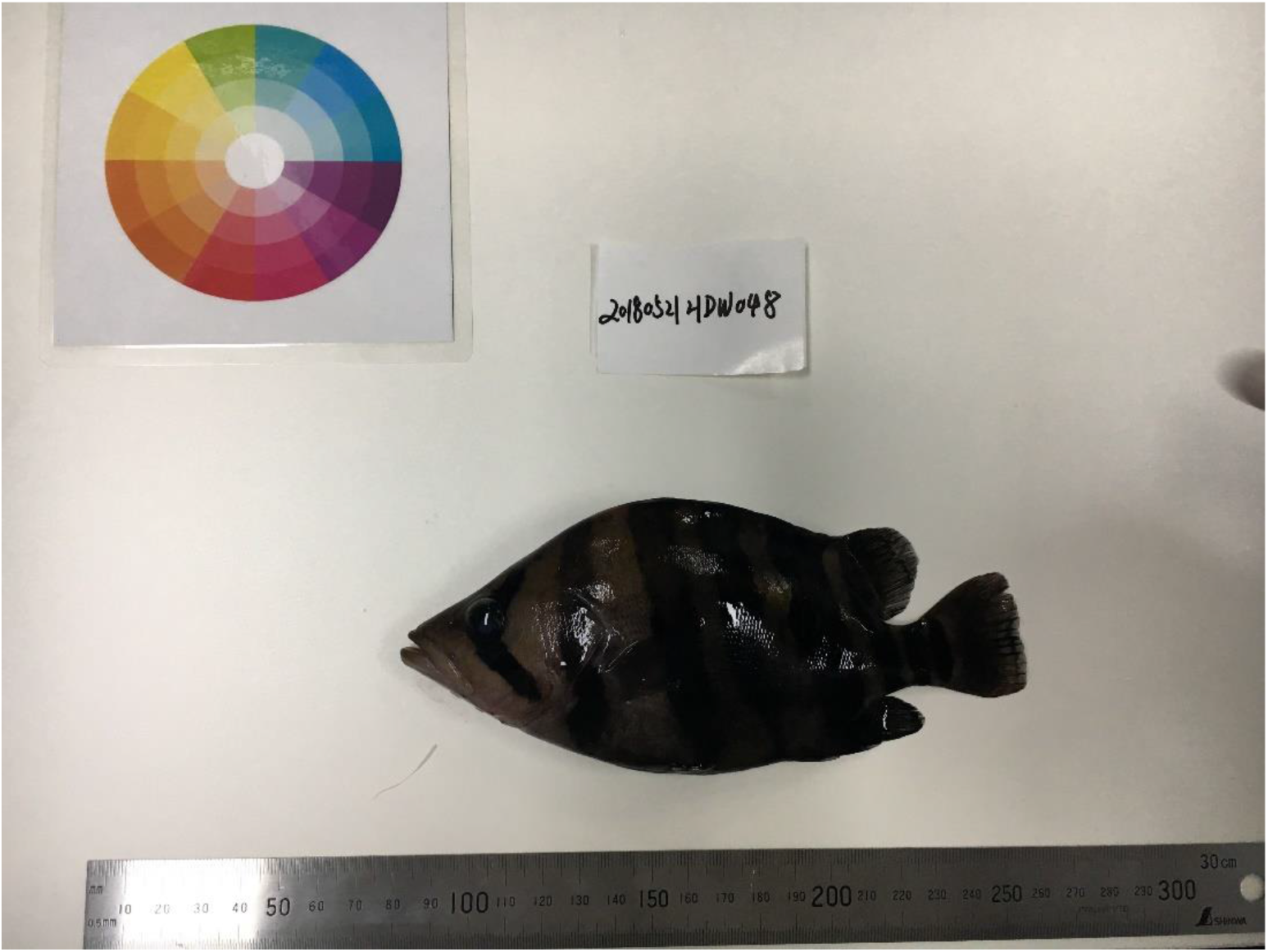
The photo of the Mekong tiger perch captured in Mekong river.

**Supplementary figure 2.**
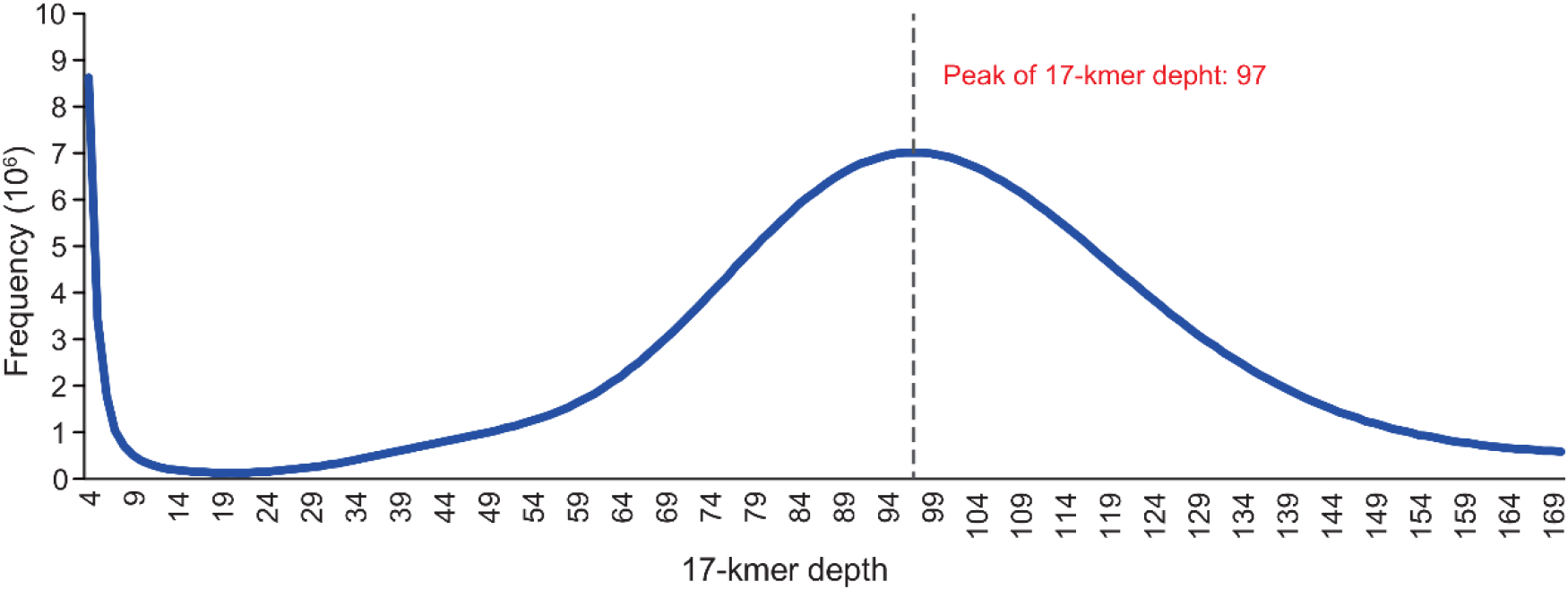
Kmer (K=17) frequency at different Kmer depth. The total number of Kmer is 60,437,782,622, the Kmer depth peak at 97, and the reads sequencing length was 100, so that the genome size was estimated 623,069,923bp.

**Supplementary figure 3.**
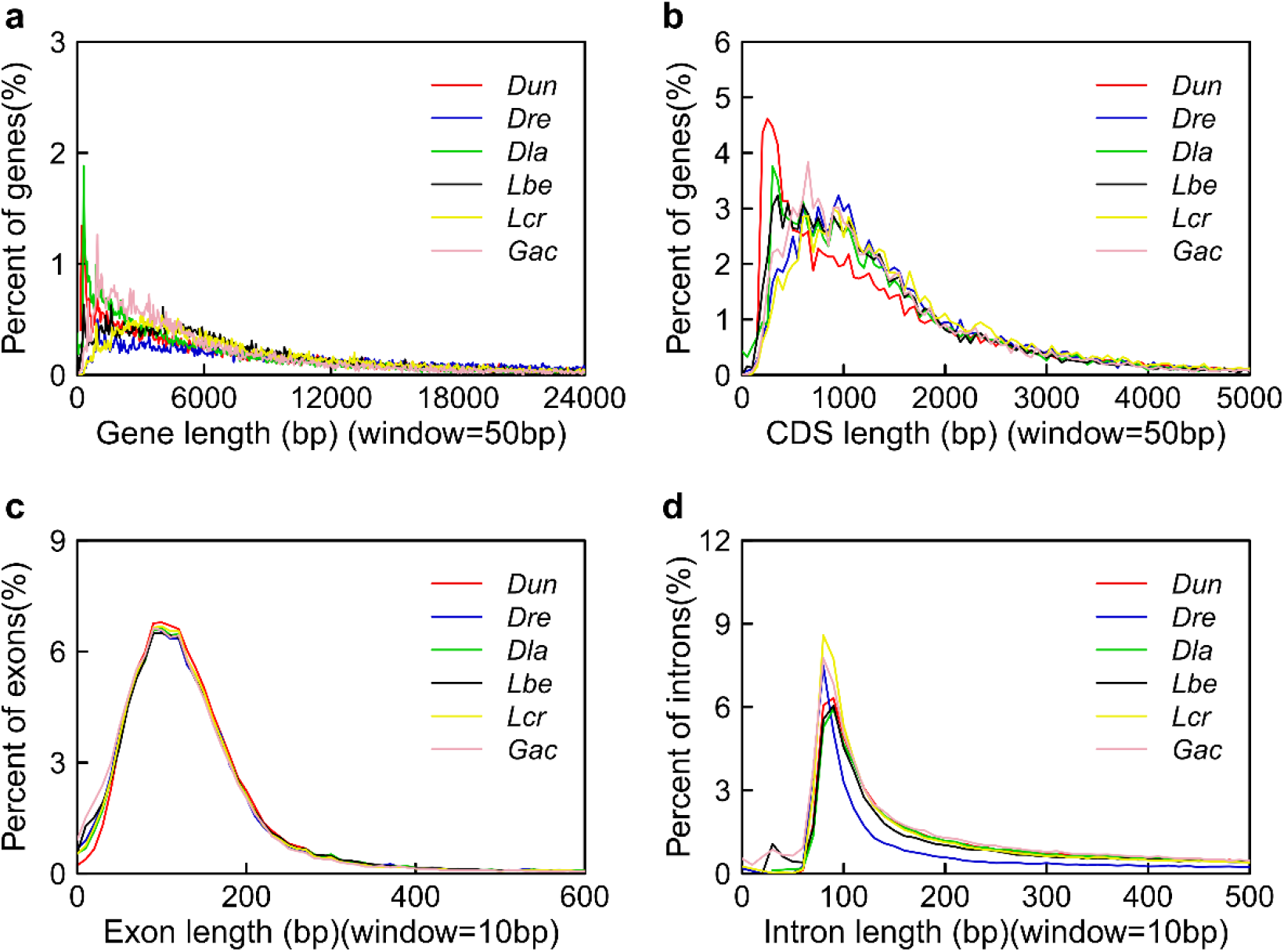
The distribution of gene length, CDS length, exon length and intron length of the Mekong tiger perch compared to related speices. Scientific names are abbreviated as follows: ***Dun***, *Datnioides undecimradiatus*; ***Dre***, *Danio rerio*; ***Dla***, *Dicentrarchus labrax*; ***Lbe***, *Labrus bergylta*; ***Lcr***, *Larimichthys crocea*; ***Gac***, *Gasterosteus aculeatus*.

**Supplementary figure 4.**
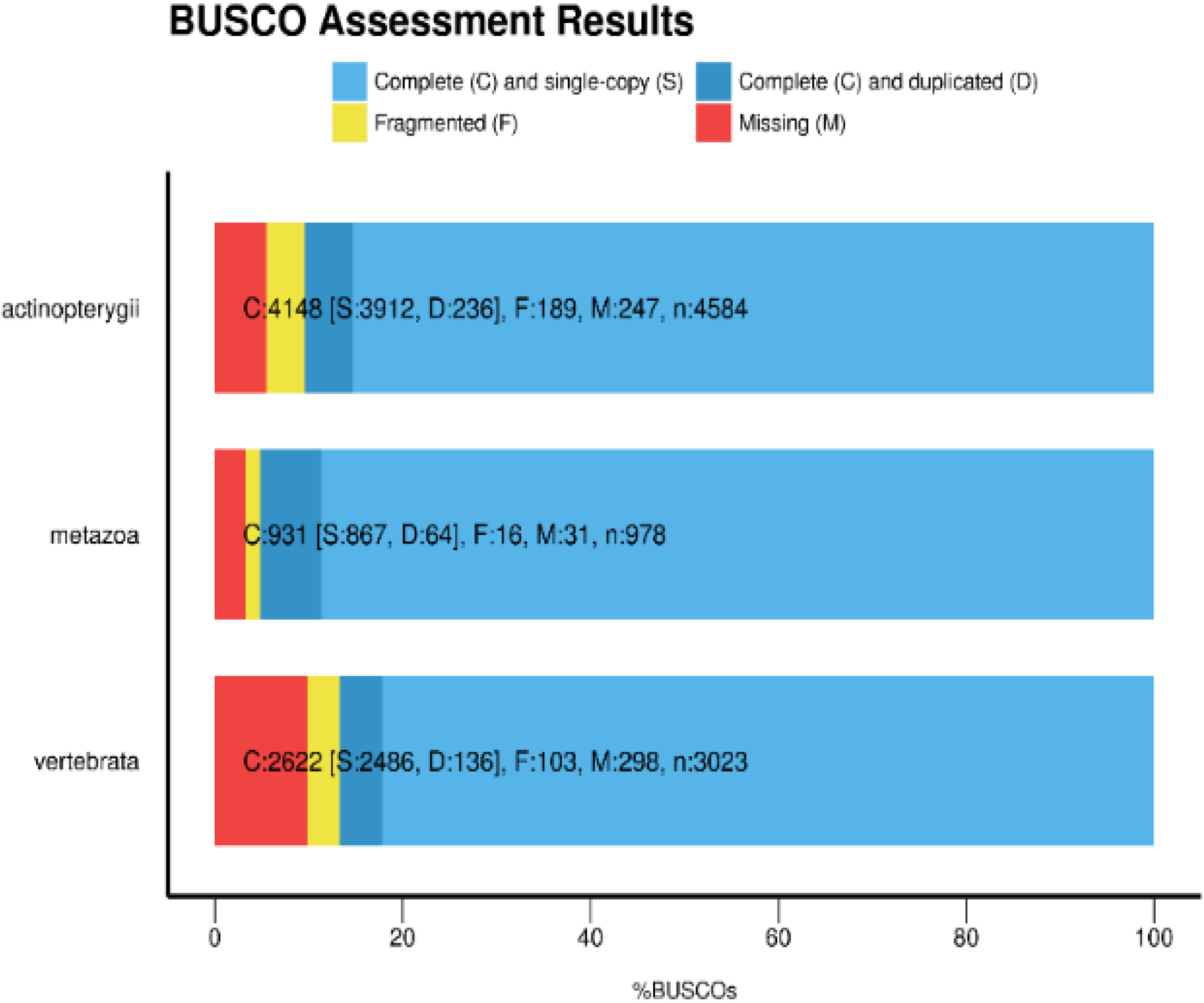
The completeness of predicted genes sets was evaluated by BUSCO based on three different databases, including Actinopterygii (v9), metazoan (v9) and vertebrata (v9).

**Supplementary figure 5.**
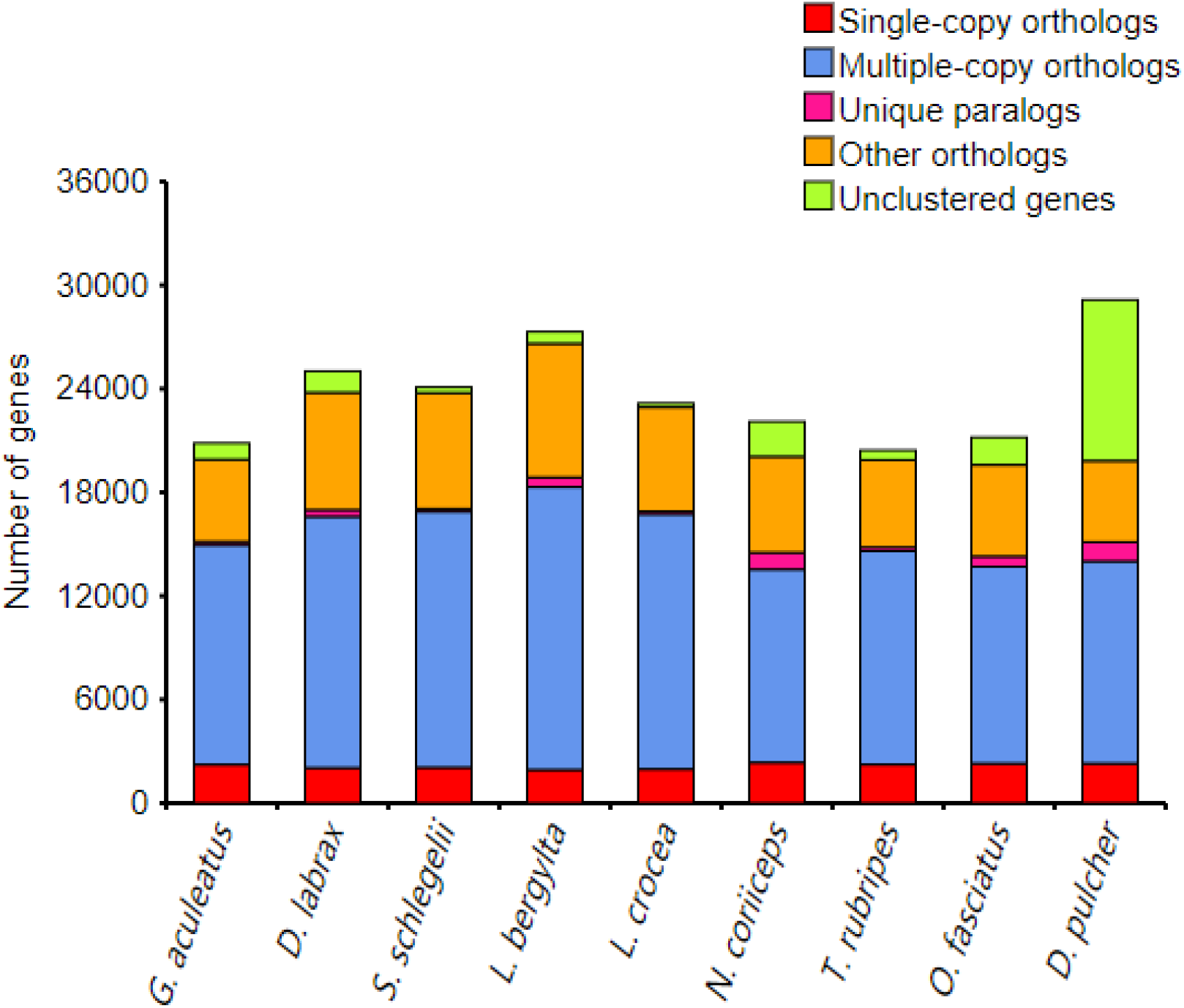
The genes number of five type of gene families for nine species used in the analysis of comparative genomes.

**Supplementary figure 6.**
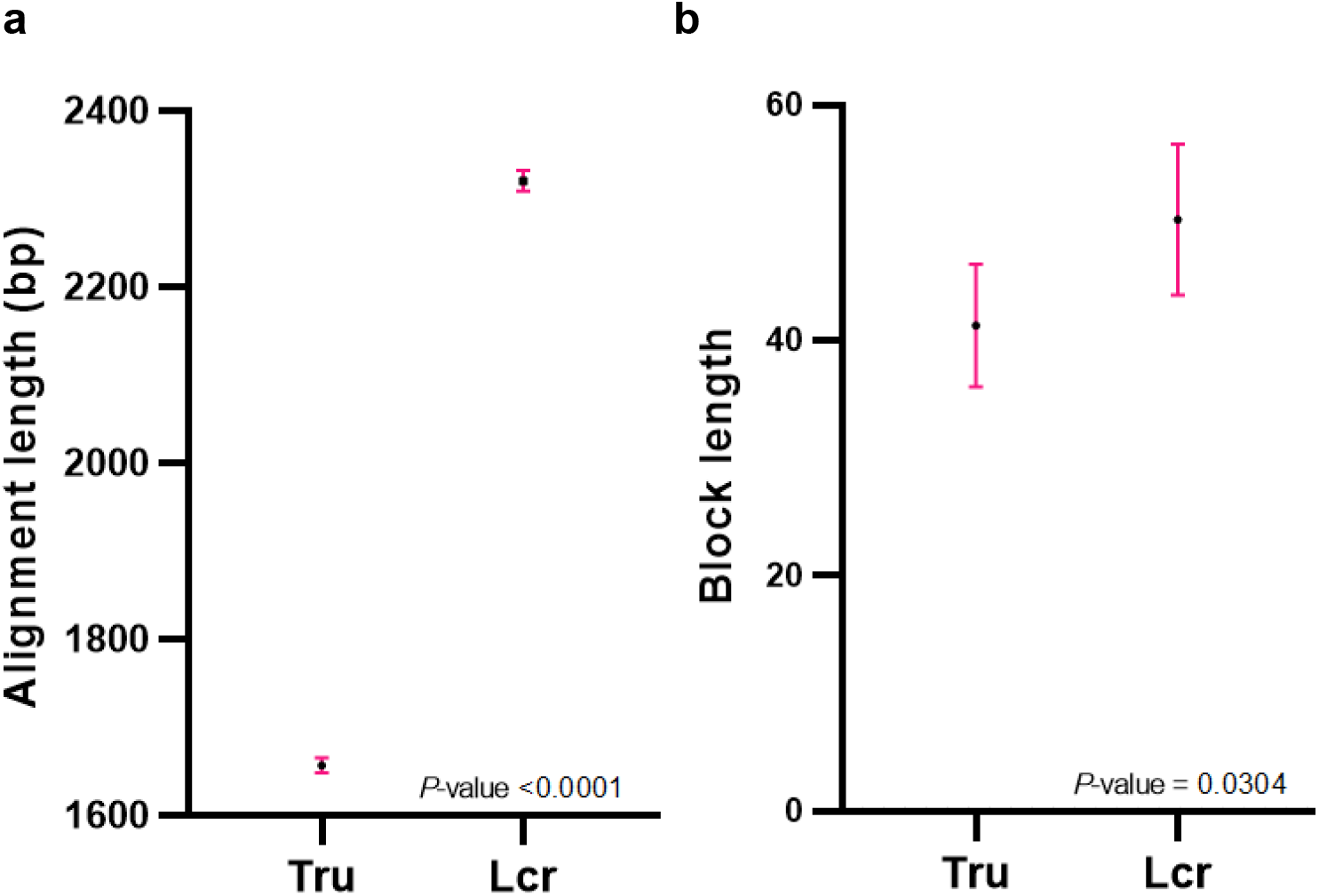
The t-test statistics of alignment length and block length. **(a)** The distribution of the length of synteny blocks at nucleotide-level. *P*-value was calculated using t-statistic. **(b)** The distribution of the length of synteny blocks at gene-level. *P*-value was using t-statistic.

**Supplementary table 1.** The details of longest 71 scaffolds.

| scaffold ID | Length (bp) | GC content (%) | N_ratio (%) |
| --- | --- | --- | --- |
| scaffold22 | 39,306,008 | 42.33 | 1.58 |
| scaffold23 | 24,901,081 | 42.33 | 1.97 |
| scaffold24 | 23,317,972 | 42.03 | 0.87 |
| scaffold63 | 22,660,142 | 42.41 | 1.76 |
| scaffold26 | 19,767,337 | 41.45 | 0.89 |
| scaffold27 | 19,537,586 | 42.15 | 1.03 |
| scaffold58 | 17,534,777 | 41.68 | 2.25 |
| scaffold25 | 17,129,097 | 41.35 | 2.02 |
| scaffold28 | 15,606,803 | 41.60 | 0.76 |
| scaffold45 | 15,041,297 | 41.93 | 1.28 |
| scaffold60 | 13,384,554 | 42.11 | 0.81 |
| scaffold61 | 13,186,598 | 42.71 | 1.04 |
| scaffold77 | 11,679,837 | 42.19 | 2.24 |
| scaffold79 | 11,352,832 | 42.26 | 0.96 |
| scaffold59 | 10,986,376 | 41.77 | 1.75 |
| scaffold88 | 10,659,222 | 41.96 | 1.16 |
| scaffold89 | 9,983,168 | 43.64 | 3.77 |
| scaffold62 | 9,730,178 | 42.24 | 1.24 |
| scaffold64 | 9,455,329 | 41.09 | 0.99 |
| scaffold80 | 9,009,409 | 41.98 | 1.13 |
| scaffold90 | 8,890,068 | 44.17 | 5.80 |
| scaffold66 | 7,986,709 | 42.51 | 0.72 |
| scaffold67 | 7,981,977 | 42.51 | 0.72 |
| scaffold155 | 7,882,590 | 44.18 | 2.10 |
| scaffold76 | 7,221,551 | 42.34 | 0.79 |
| scaffold83 | 7,024,340 | 43.41 | 1.36 |
| scaffold129 | 7,004,772 | 43.25 | 3.18 |
| scaffold86 | 6,951,544 | 44.19 | 2.53 |
| scaffold87 | 6,939,792 | 42.63 | 1.33 |
| scaffold85 | 6,732,700 | 43.19 | 2.76 |
| scaffold75 | 6,668,441 | 40.74 | 0.60 |
| scaffold102 | 6,529,773 | 42.51 | 1.49 |
| scaffold78 | 6,505,070 | 40.03 | 1.04 |
| scaffold110 | 6,218,238 | 42.57 | 2.73 |
| scaffold65 | 6,149,668 | 41.30 | 0.92 |
| scaffold81 | 6,042,282 | 40.32 | 0.84 |
| scaffold107 | 5,552,807 | 43.61 | 1.57 |
| scaffold82 | 5,383,449 | 45.41 | 3.42 |
| scaffold108 | 5,037,774 | 42.28 | 1.01 |
| scaffold111 | 4,837,472 | 43.64 | 1.99 |
| scaffold105 | 4,507,353 | 42.91 | 1.23 |
| scaffold101 | 4,379,686 | 46.21 | 5.84 |
| scaffold176 | 4,245,196 | 46.52 | 3.11 |
| scaffold125 | 4,183,443 | 46.18 | 1.54 |
| scaffold84 | 3,961,987 | 40.27 | 0.99 |
| scaffold106 | 3,649,163 | 43.70 | 1.26 |
| scaffold148 | 3,238,164 | 41.03 | 2.37 |
| scaffold147 | 3,070,712 | 47.24 | 2.74 |
| scaffold103 | 2,911,789 | 45.84 | 9.75 |
| scaffold69 | 2,816,955 | 45.17 | 5.65 |
| scaffold68 | 2,814,068 | 45.16 | 5.64 |
| scaffold131 | 2,714,138 | 42.91 | 2.44 |
| scaffold186 | 2,711,499 | 39.58 | 0.54 |
| scaffold200 | 2,366,470 | 46.18 | 4.82 |
| scaffold113 | 2,352,837 | 42.56 | 3.55 |
| scaffold104 | 2,165,782 | 41.93 | 0.79 |
| scaffold201 | 2,162,358 | 43.22 | 0.86 |
| scaffold122 | 2,118,146 | 46.78 | 8.16 |
| scaffold133 | 2,111,711 | 41.92 | 4.03 |
| scaffold18494 | 1,965,245 | 45.72 | 4.53 |
| scaffold18500 | 1,963,021 | 45.71 | 4.53 |
| scaffold124 | 1,863,279 | 46.42 | 5.63 |
| scaffold112 | 1,679,604 | 41.45 | 0.25 |
| scaffold109 | 1,675,575 | 42.10 | 2.24 |
| scaffold137 | 1,624,607 | 42.65 | 3.82 |
| scaffold173 | 1,554,608 | 43.17 | 0.68 |
| scaffold130 | 1,511,149 | 41.63 | 2.51 |
| scaffold185 | 1,502,512 | 43.66 | 3.3 |
| scaffold136 | 1,485,443 | 43.24 | 2.61 |
| scaffold127 | 1,411,413 | 46.76 | 6.74 |
| scaffold135 | 1,407,639 | 47.20 | 6.86 |

**Supplementary table 2.** Repeat annotation of the Mekong tiger perch genome.

| Type | Rebase TEs |  | TE proteins |  | De novo |  | Combined TEs |  |
| --- | --- | --- | --- | --- | --- | --- | --- | --- |
|  | Length (bp) | % in genome | Length (bp) | % in genome | Length (bp) | % in genome | Length (bp) | % in genome |
| DNA | 16,506,661 | 2.77 | 1,633,161 | 0.27 | 30,824,896 | 5.18 | 36,324,964 | 6.11 |
| LINE | 6,821,950 | 1.15 | 4,071,954 | 0.68 | 12,961,236 | 2.18 | 16,349,136 | 2.75 |
| SINE | 662,765 | 0.11 | - | 0.00 | 908,953 | 0.15 | 1,152,466 | 0.19 |
| LTR | 6,332,971 | 1.06 | 1,837,976 | 0.31 | 9,602,730 | 1.61 | 14,084,966 | 2.37 |
| Other | 4,319 | 0.00 | - | 0.00 | - | 0.00 | 4,319 | 0.00 |
| Unknown | - | 0.00 | - | 0.00 | 19,939,915 | 3.35 | 19,939,915 | 3.35 |
| Total | 26,352,522 | 4.43 | 7,536,991 | 1.27 | 65,356,559 | 10.98 | 71,231,464 | 11.97 |

**Supplementary table 3.** The statistics of predicted genes using different methods.

| Method | Software/Species | Number of<br>predicted genes | Average gene<br>length (bp) | Average CDS<br>length (bp) | Average exon<br>number | Average exon<br>length (bp) | Average intron<br>length (bp) |
| --- | --- | --- | --- | --- | --- | --- | --- |
| *ab initio* | Augustus | 26,826 | 11324.91 | 1428.76 | 8.21 | 173.94 | 1371.75 |
|  | Genscan | 29,524 | 14516.5 | 1596.21 | 9.18 | 173.87 | 1579.41 |
| Homolog-based | *Danio rerio* | 44,300 | 19786.39 | 1526.41 | 8.79 | 173.62 | 2343.59 |
|  | *Dicentrarchus labrax* | 28,264 | 12505.75 | 1483.79 | 8.13 | 182.44 | 1545.24 |
|  | *Labrus bergylta* | 39,412 | 21851.38 | 1494.87 | 8.67 | 172.36 | 2653.06 |
|  | *Larimichthys crocea* | 47,139 | 19654.28 | 2187.95 | 12.62 | 173.42 | 1503.63 |
|  | *Gasterosteus aculeatus* | 28,975 | 10277.3 | 1456.49 | 8.92 | 163.2 | 1113.07 |
| Combined | EVM | 29,150 | 13758.59 | 1510.42 | 9.04 | 167.03 | 1522.83 |

**Supplementary table 4.**
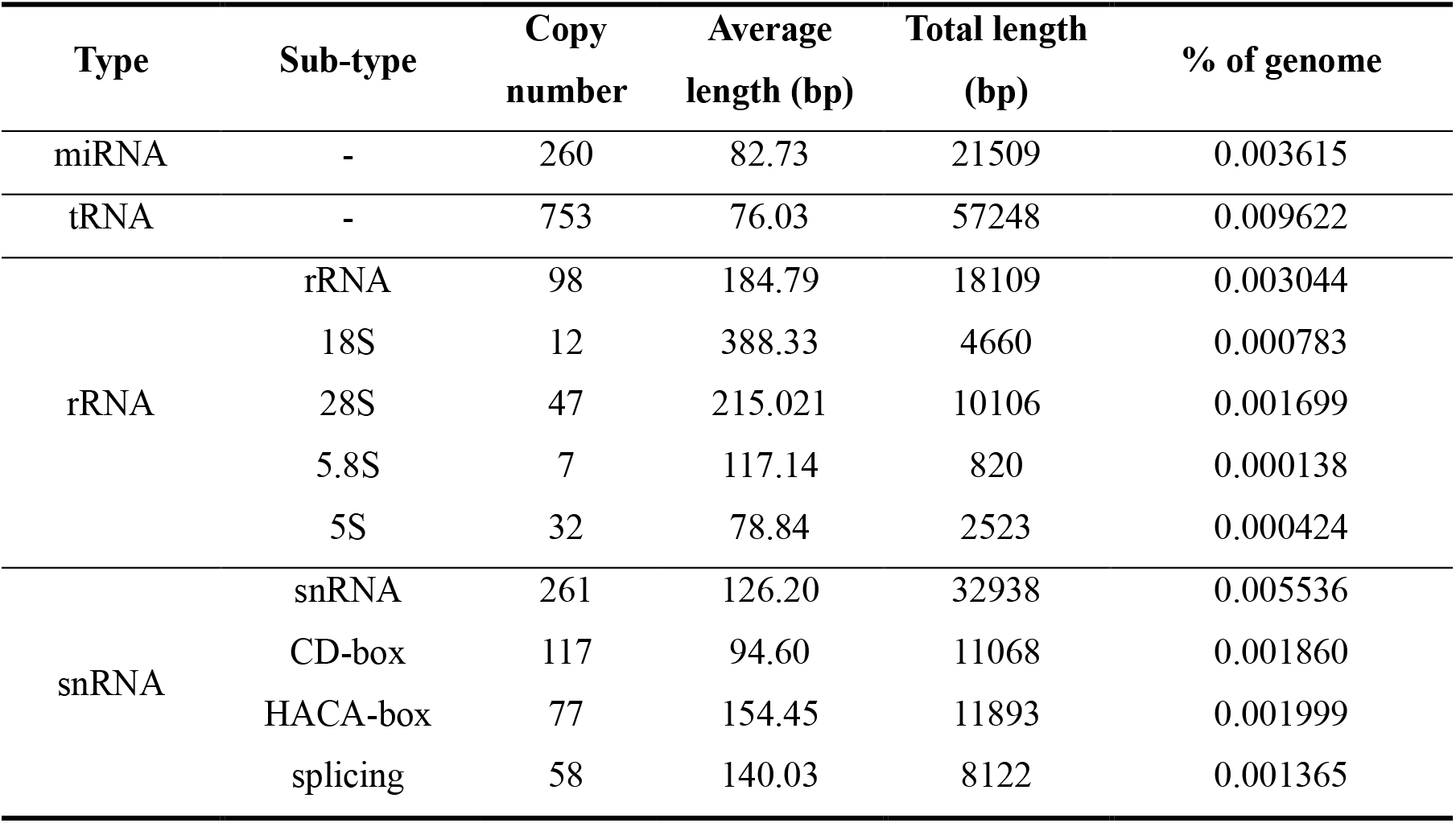
ncRNA annotation of the Mekong tiger perch genome.

| Type | Sub-type | Copy number | Average length (bp) | Total length (bp) | % of genome |
| --- | --- | --- | --- | --- | --- |
| miRNA | - | 260 | 82.73 | 21509 | 0.003615 |
| tRNA | - | 753 | 76.03 | 57248 | 0.009622 |
| rRNA | rRNA | 98 | 184.79 | 18109 | 0.003044 |
|  | 18S | 12 | 388.33 | 4660 | 0.000783 |
|  | 28S | 47 | 215.021 | 10106 | 0.001699 |
|  | 5.8S | 7 | 117.14 | 820 | 0.000138 |
|  | 5S | 32 | 78.84 | 2523 | 0.000424 |
| snRNA | snRNA | 261 | 126.20 | 32938 | 0.005536 |
|  | CD-box | 117 | 94.60 | 11068 | 0.001860 |
|  | HACA-box | 77 | 154.45 | 11893 | 0.001999 |
|  | splicing | 58 | 140.03 | 8122 | 0.001365 |

**Supplementary table 5.** Genes annotated on mitochondrial genome.

| Mitochondrial genome | Start | End | Length(bp) | Direction | Type | Gene name | Gene product | Occurred Counts |
| --- | --- | --- | --- | --- | --- | --- | --- | --- |
| C515351 | 134 | 202 | 69 | + | tRNA | trnF(gaa) | tRNA-Phe | 1 |
| C515351 | 202 | 1170 | 969 | + | rRNA | s-rRNA | 12S ribosomal RNA | 1 |
| C515351 | 1170 | 1242 | 73 | + | tRNA | trnV(uac) | tRNA-Val | 1 |
| C515351 | 1262 | 2954 | 1693 | + | rRNA | l-rRNA | 16S ribosomal RNA | 1 |
| C515351 | 2954 | 3028 | 75 | + | tRNA | trnL(uaa) | tRNA-Leu | 2 |
| C515351 | 3028 | 4003 | 976 | + | CDS | ND1 | NADH dehydrogenase subunit 1 | 1 |
| C515351 | 4008 | 4079 | 72 | + | tRNA | trnI(gau) | tRNA-Ile | 1 |
| C515351 | 4078 | 4149 | 72 | - | tRNA | trnQ(uug) | tRNA-Gln | 1 |
| C515351 | 4148 | 4220 | 73 | + | tRNA | trnM(cau) | tRNA-Met | 1 |
| C515351 | 4220 | 5267 | 1048 | + | CDS | ND2 | NADH dehydrogenase subunit 2 | 1 |
| C515351 | 5266 | 5338 | 73 | + | tRNA | trnW(uca) | tRNA-Trp | 1 |
| C515351 | 5338 | 5407 | 70 | - | tRNA | trnA(ugc) | tRNA-Ala | 1 |
| C515351 | 5408 | 5481 | 74 | - | tRNA | trnN(guu) | tRNA-Asn | 1 |
| C515351 | 5516 | 5585 | 70 | - | tRNA | trnC(gca) | tRNA-Cys | 1 |
| C515351 | 5585 | 5655 | 71 | - | tRNA | trnY(gua) | tRNA-Tyr | 1 |
| C515351 | 5656 | 7207 | 1552 | + | CDS | COX1 | cytochrome c oxidase subunit I | 1 |
| C515351 | 7207 | 7278 | 72 | - | tRNA | trnS(uga) | tRNA-Ser | 2 |
| C515351 | 7281 | 7354 | 74 | + | tRNA | trnD(guc) | tRNA-Asp | 1 |
| C515351 | 7362 | 8061 | 700 | + | CDS | COX2 | cytochrome c oxidase subunit II | 1 |
| C515351 | 8053 | 8128 | 76 | + | tRNA | trnK(uuu) | tRNA-Lys | 1 |
| C515351 | 8129 | 8297 | 169 | + | CDS | ATP8 | ATP synthase F0 subunit 8 | 1 |
| C515351 | 8314 | 8971 | 658 | + | CDS | ATP6 | ATP synthase F0 subunit 6 | 1 |
| C515351 | 8970 | 9756 | 787 | + | CDS | COX3 | cytochrome c oxidase subunit III | 1 |
| C515351 | 9755 | 9827 | 73 | + | tRNA | trnG(ucc) | tRNA-Gly | 1 |
| C515351 | 9827 | 10178 | 352 | + | CDS | ND3 | NADH dehydrogenase subunit 3 | 1 |
| C515351 | 10176 | 10245 | 70 | + | tRNA | trnR(ucg) | tRNA-Arg | 1 |
| C515351 | 10245 | 10542 | 298 | + | CDS | ND4L | NADH dehydrogenase 4L | 1 |
| C515351 | 10535 | 11921 | 1387 | + | CDS | ND4 | NADH dehydrogenase 4 | 1 |
| C515351 | 11916 | 11984 | 69 | + | tRNA | trnH(gug) | tRNA-His | 1 |
| C515351 | 11984 | 12051 | 68 | + | tRNA | trnS(gcu) | tRNA-Ser | 2 |
| C515351 | 12055 | 12128 | 74 | + | tRNA | trnL(uag) | tRNA-Leu | 2 |
| C515351 | 12128 | 13967 | 1840 | + | CDS | ND5 | NADH dehydrogenase subunit 5 | 1 |
| C515351 | 13963 | 14485 | 523 | - | CDS | ND6 | NADH dehydrogenase 6 | 1 |
| C515351 | 14486 | 14555 | 70 | - | tRNA | trnE(uuc) | tRNA-Glu | 1 |
| C515351 | 14560 | 15721 | 1162 | + | CDS | CYTb | cytochrome b | 1 |
| C515351 | 15701 | 15771 | 71 | + | tRNA | trnT(ugu) | tRNA-Thr | 1 |
| C515351 | 15770 | 15840 | 71 | - | tRNA | trnP(ugg) | tRNA-Pro | 1 |

**Supplementary table 6.** Relationship GC with repeat content.

| Pairs | Pearson r | Lower 95% CI | Upper 95% CI | P-value |
| --- | --- | --- | --- | --- |
| CGI density vs. GC content | 0.643 | 0.666 | 0.688 | 4.83E-304 |
| CGI density vs. Gene density | 0.252 | 0.290 | 0.326 | 4.37E-47 |
| CGI density vs. Repeat content | 0.329 | 0.364 | 0.399 | 1.75E-75 |
| GC content vs. Gene density | 0.135 | 0.174 | 0.213 | 1.42E-17 |
| GC content vs. Repeat content | 0.108 | 0.147 | 0.187 | 5.27E-13 |
| Gene density vs. Repeat content | -0.082 | -0.042 | -0.002 | 4.16E-02 |

**Supplementary table 7.** Nine species used in our study.

| Three letter code | Scientific name | Common name | Order | Data source | Accession ID |
| --- | --- | --- | --- | --- | --- |
| *Ofa* | *Oplegnathus fasciatus* | barred knifejaw | Centrarchiformes | NCBI | GCA_003416845.1 |
| *Lbe* | *Labrus bergylta* | ballan wrasse | Labriiformes | NCBI | GCF_900080235.1 |
| *Dun* | *Datnioides undecimradiatus* | Mekong tiger perch | Lobotiformes | our study | our study |
| *Dla* | *Dicentrarchus labrax* | European seabass | Moronidae | NCBI | GCA_000689215.1 |
| *Gac* | *Gasterosteus aculeatus* | three-spined stickleback | Perciformes | NCBI | GCA_000180675.1 |
| *Nco* | *Notothenia coriiceps* | black rockcod | Perciformes | NCBI | GCF_000735185.1 |
| *Ssc* | *Sebastes schlegelii* | Schlegel's black rockfish | Perciformes | NCBI | GCA_004335315.1 |
| *Lcr* | *Larimichthys crocea* | large yellow croaker | Sciaenidae | NCBI | GCF_000972845.2 |
| *Tru* | *Takifugu rubripes* | torafugu | Tetraodontiformes | NCBI | GCF_000180615.1 |

**Supplementary table 8.** The statistics of gene family clusters.

| Spec<br>ies | Genes*<br>number | Un-clustered<br>genes number | Families<br>number | Unique family's<br>number | Average genes<br>per family |
| --- | --- | --- | --- | --- | --- |
| *Gac* | 20,862 | 1,107 | 8,792 | 19 | 2.25 |
| *Dla* | 25,020 | 1,423 | 10,090 | 77 | 2.34 |
| *Ssc* | 24,094 | 425 | 9,745 | 21 | 2.43 |
| *Lbe* | 27,305 | 888 | 10,031 | 133 | 2.63 |
| *Lcr* | 23,163 | 315 | 9,967 | 30 | 2.29 |
| *Nco* | 22,099 | 2,713 | 9,752 | 139 | 1.99 |
| *Tru* | 20,434 | 704 | 9,359 | 37 | 2.11 |
| *Ofa* | 21,901 | 2,023 | 9,481 | 81 | 2.10 |
| *Dun* | 26,894 | 8,260 | 9,208 | 378 | 2.02 |

\* Pseudo-genes (stop-gained in middle) were filtered out, and the genes with protein length <50 were also removed.

**Supplementary table 9.** Synteny of alignment statistics at whole-genome nucleotide sequences level.

| Species aligned to<br>*Dun* | Cutoff of alignment length | Synteny coverage (%) | Alignment number | Average alignment length (bp) | Median alignment length (bp) | Min alignment length (bp) | Max alignment length (bp) |
| --- | --- | --- | --- | --- | --- | --- | --- |
| *Lcr*<br>(length:658 M) | ≥ 1kb | 41.13% | 105,447 | 2,321 | 1,717 | 1,000 | 56,540 |
|  | ≥ 2kb | 26.02% | 41,422 | 3,738 | 2,967 | 2,000 | 56,540 |
|  | ≥ 5kb | 8.89% | 6,811 | 7,769 | 6,571 | 5,000 | 56,540 |
| *Tru*<br>(length:391 M) | ≥ 1kb | 10.04% | 36,032 | 1,657 | 1,388 | 1,000 | 24,987 |
|  | ≥ 2kb | 3.58% | 7,312 | 2,552 | 2,954 | 2,000 | 24,987 |
|  | ≥ 5kb | 0.39% | 352 | 6,621 | 5,952 | 5,000 | 24,987 |

**Supplementary table 10.**
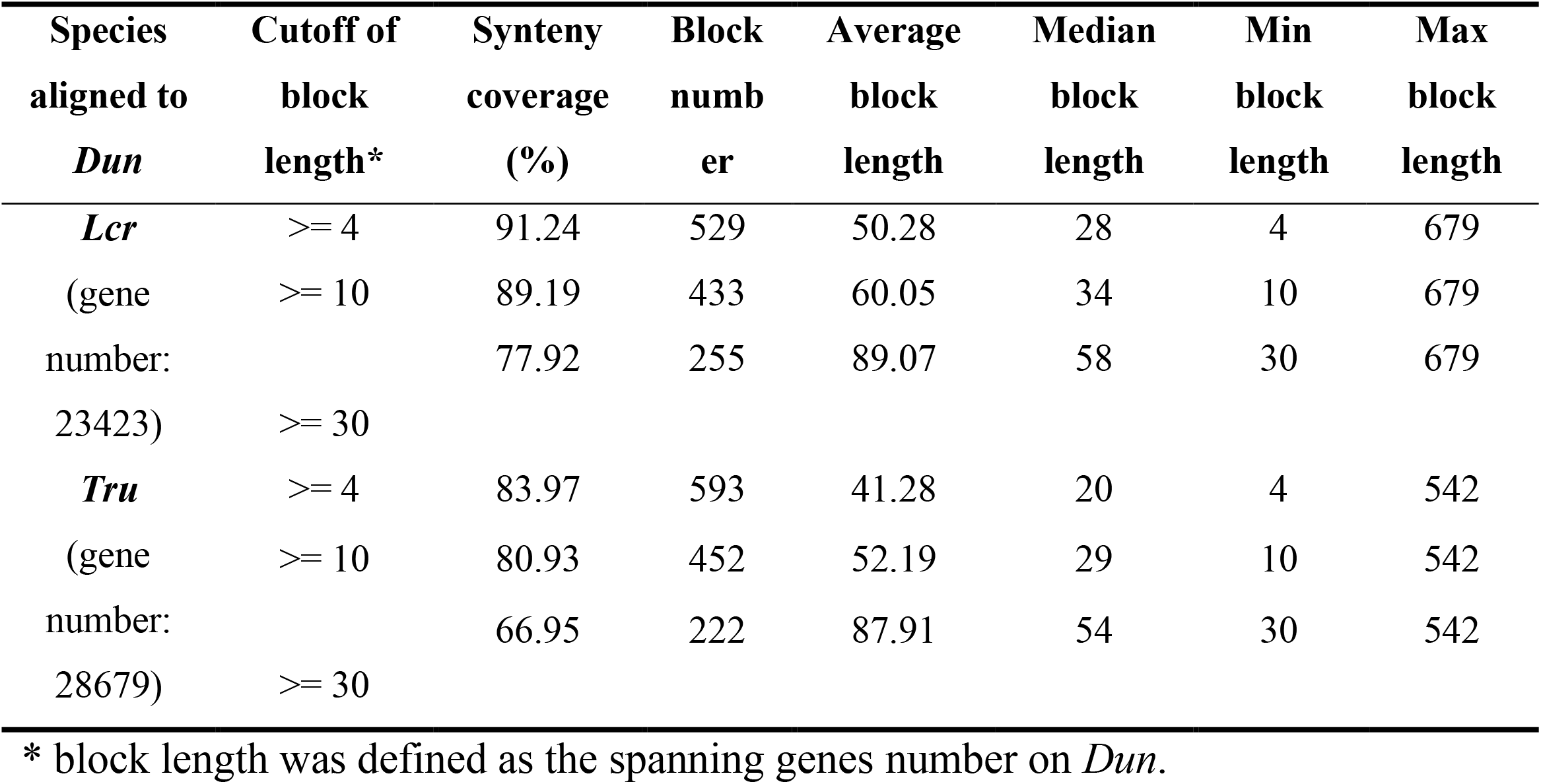
Distribution of synteny of alignment statistics at whole genes level at different cutoff.

**Supplementary table 11.** Genome assembly and gene set quality by BUSCO.

|  |  | Genome level |  | Gene level |  |
| --- | --- | --- | --- | --- | --- |
|  |  | *Larimichthys crocea* | *Takifugu rubripes* | *Larimichthys crocea* | *Takifugu rubripes* |
| actinopterygii (db_v9) | Complete BUSCOs (C) | 4362 (95.2%) | 4263 (93.0%) | 4554 (99.3%) | 4345 (94.7%) |
|  | Complete and single-copy BUSCOs (S) | 4275 (93.3%) | 4168 (90.9%) | 2615 (57.0%) | 3000 (65.4%) |
|  | Complete and duplicated BUSCOs (D) | 87 (1.9%) | 95 (2.1%) | 1939 (42.3%) | 1345 (29.3%) |
|  | Fragmented BUSCOs (F) | 119 (2.6%) | 202 (4.4%) | 23 (0.5%) | 134 (2.9%) |
|  | Missing BUSCOs (M) | 103 (2.2%) | 119 (2.6%) | 7 (0.2%) | 105 (2.4%) |
|  | Total BUSCO groups searched | 4584 | 4584 | 4584 | 4584 |
| metazoa (db_v9) | Complete BUSCOs (C) | 918 (93.9%) | 915 (93.6%) | 973 (99.5%) | 950 (97.1%) |
|  | Complete and single-copy BUSCOs (S) | 882 (90.2%) | 880 (90.0%) | 656 (67.1%) | 670 (68.5%) |
|  | Complete and duplicated BUSCOs (D) | 36 (3.7%) | 35 (3.6%) | 317 (32.4%) | 280 (28.6%) |
|  | Fragmented BUSCOs (F) | 7 (0.7%) | 12 (1.2%) | 4 (0.4%) | 15 (1.5%) |
|  | Missing BUSCOs (M) | 53 (5.4%) | 51 (5.2%) | 1 (0.1%) | 13 (1.4%) |
|  | Total BUSCO groups searched | 978 | 978 | 978 | 978 |

**Supplementary table 12.**
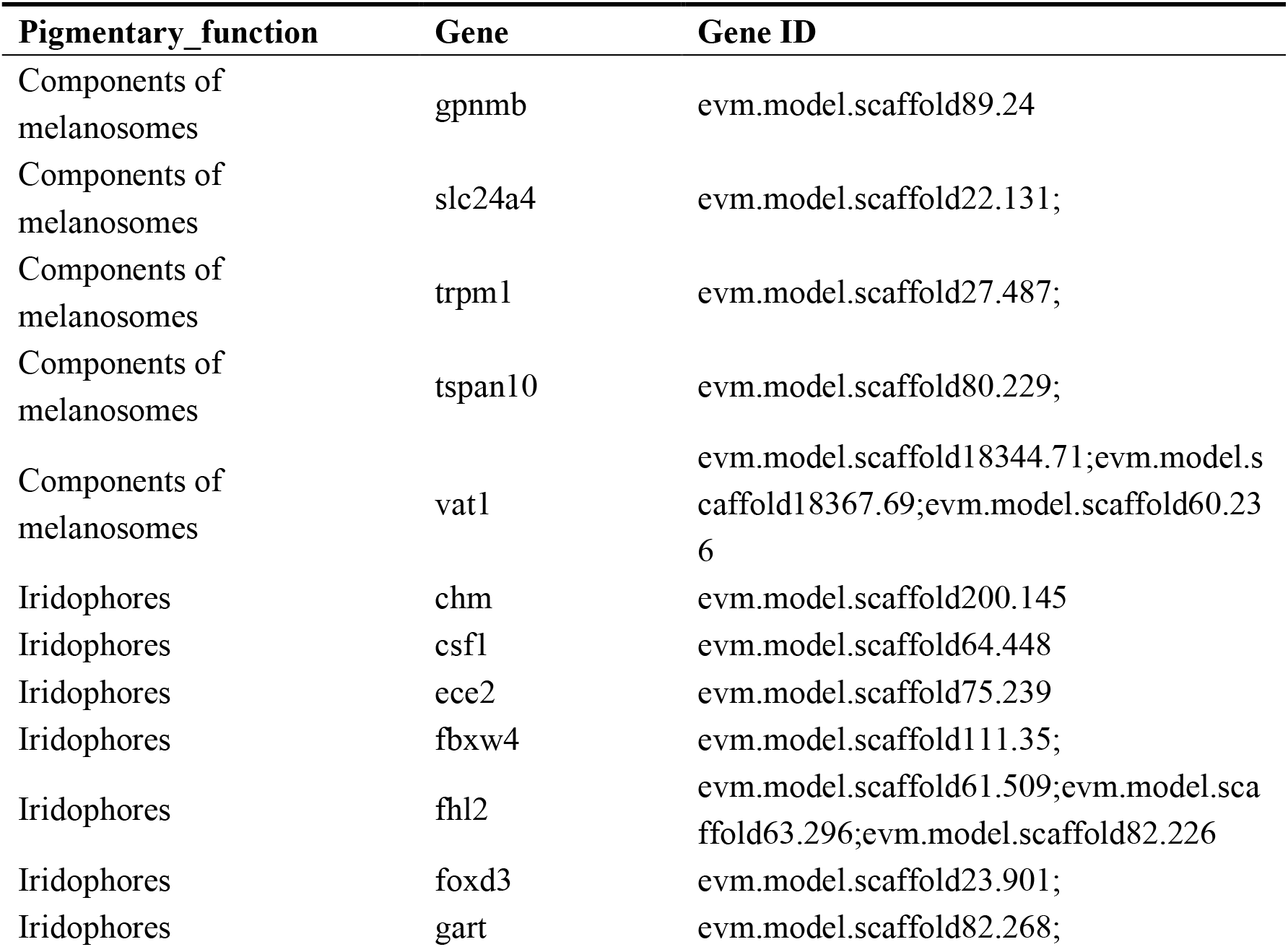
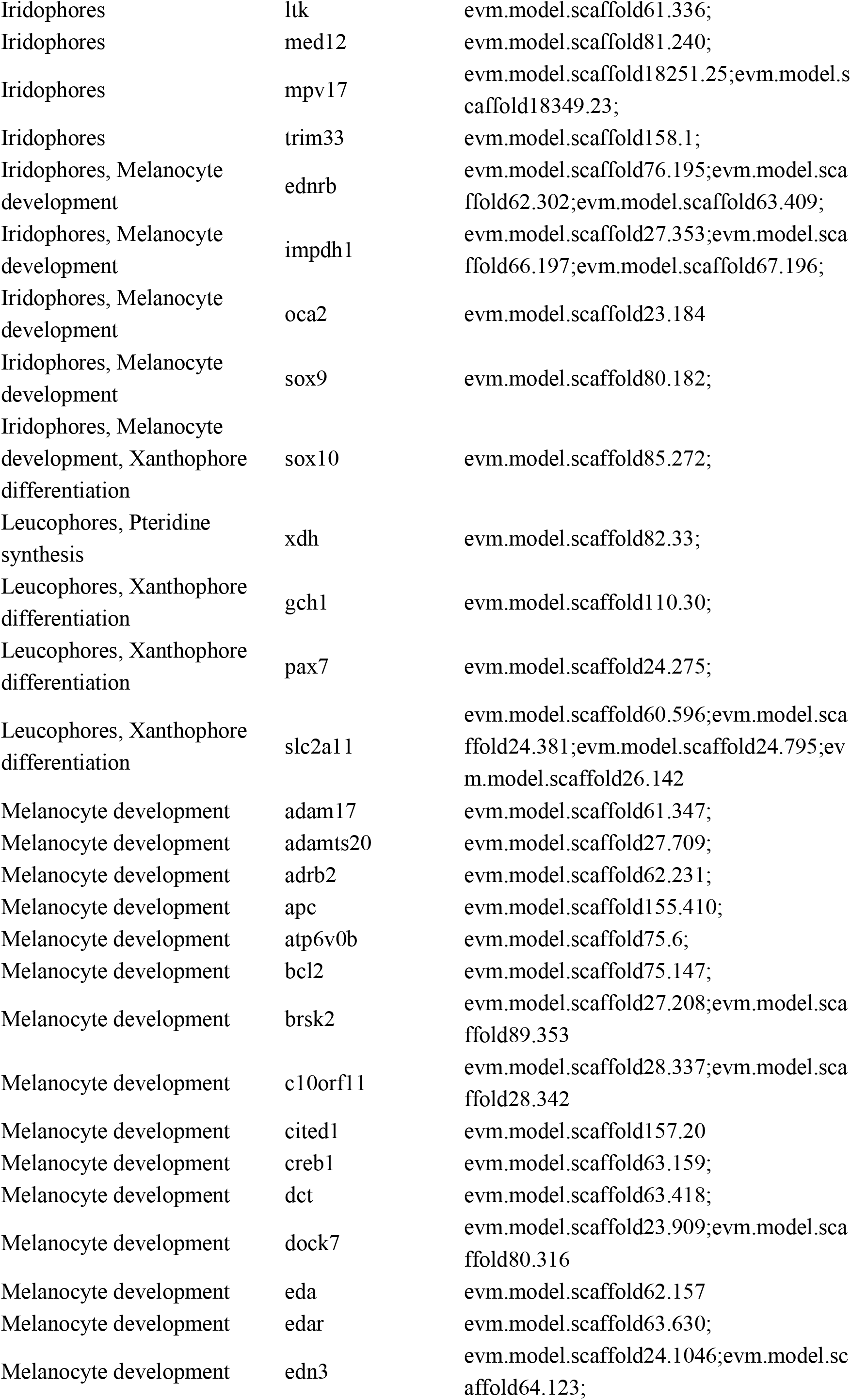
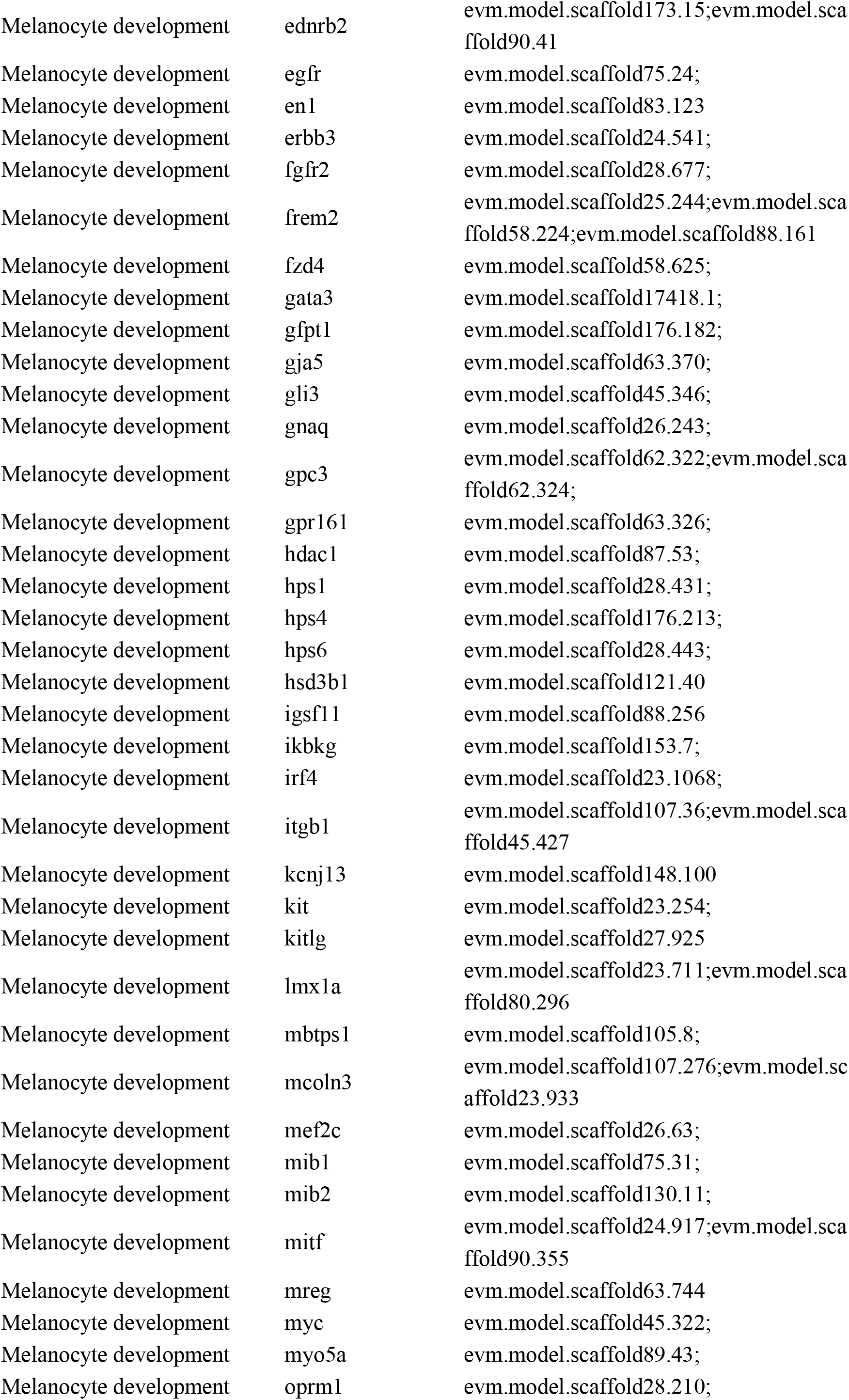
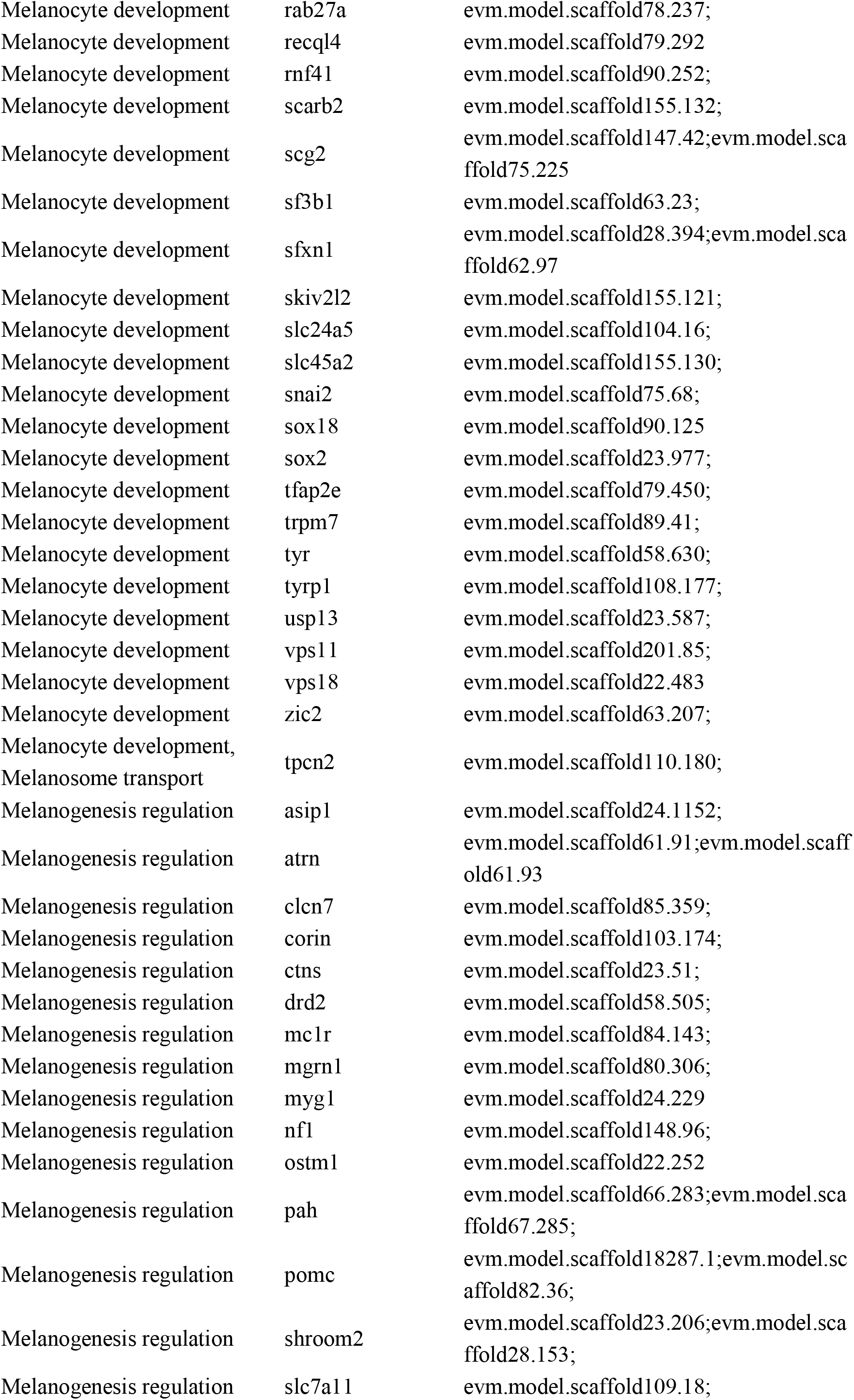
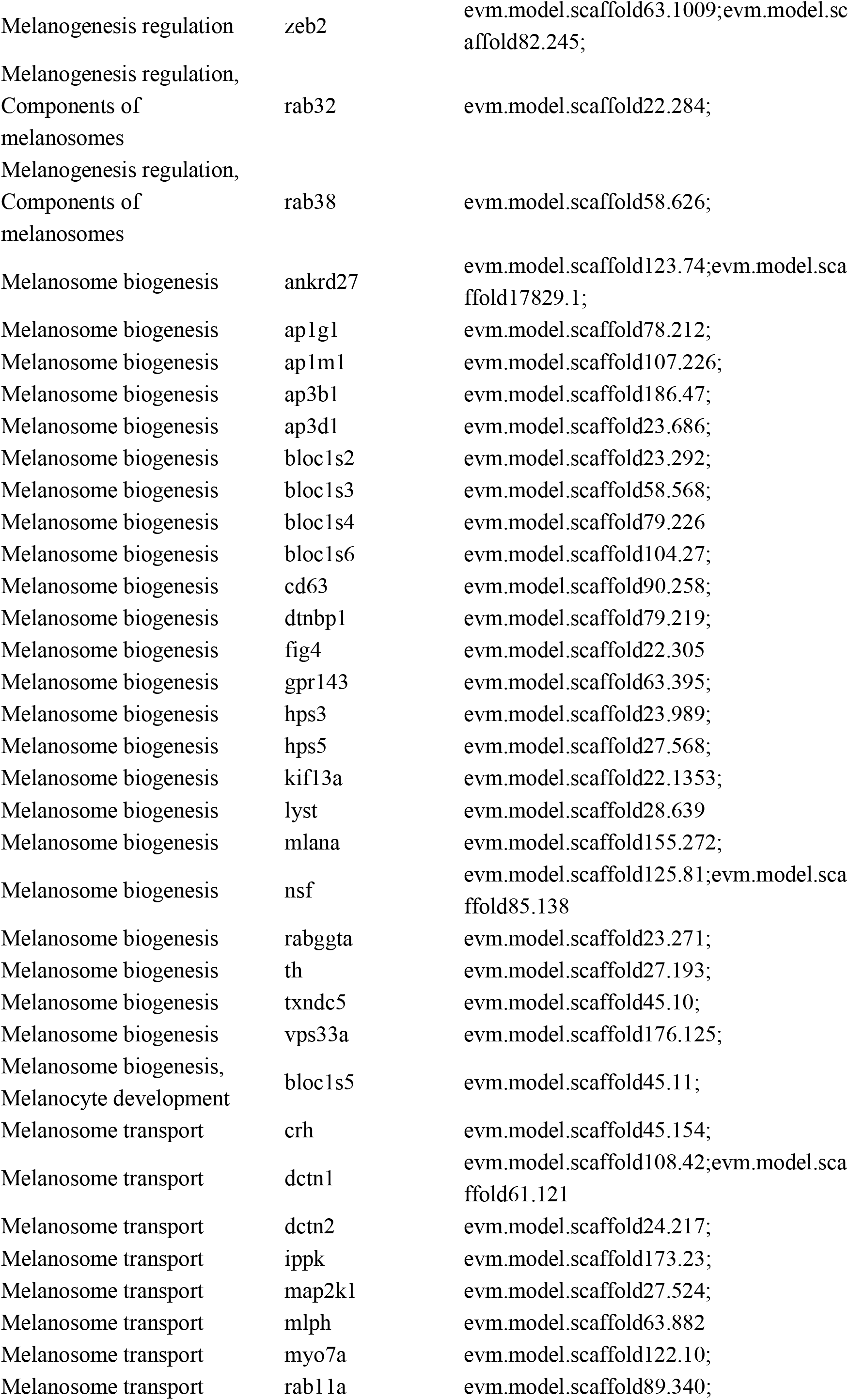
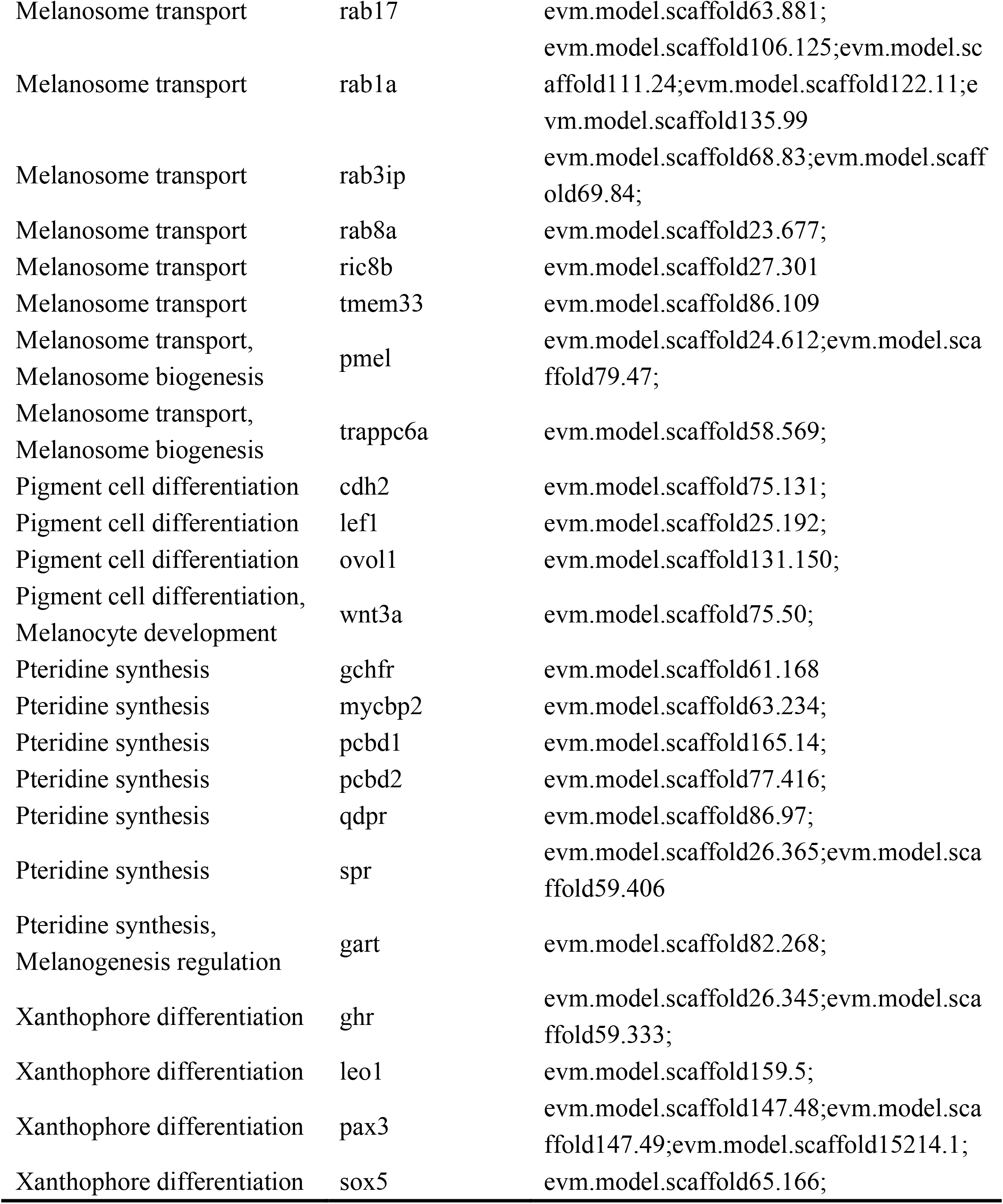
Genes involved in the pigment synthesis.

**Supplementary table 13.**
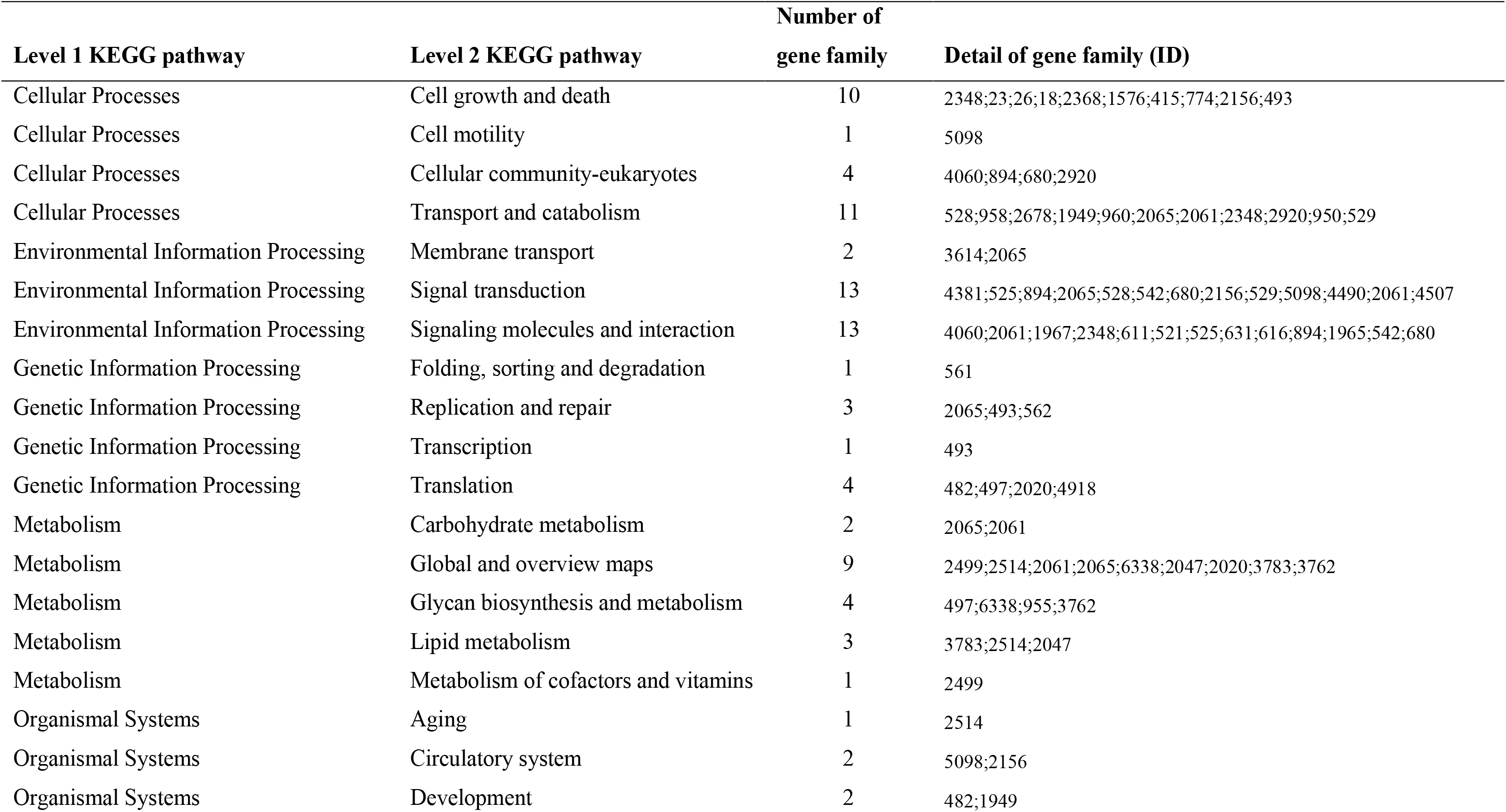
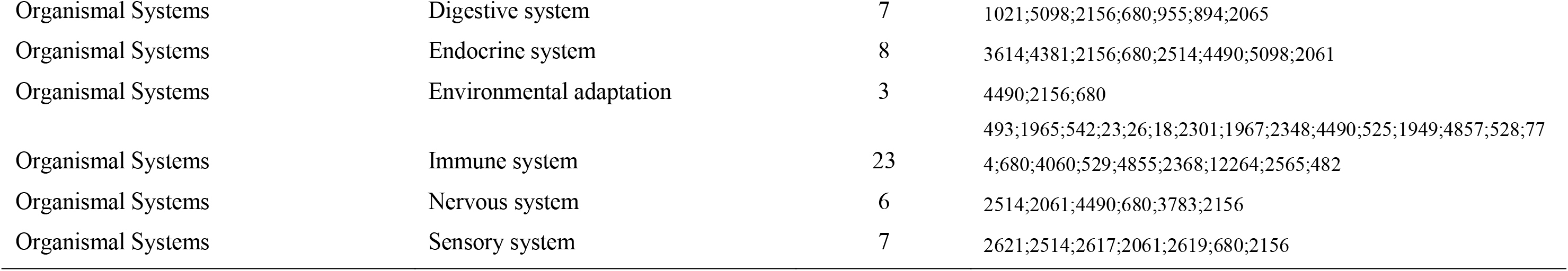
KEGG pathway involved by contracted gene families

**Supplementary table 14.** KEGG pathway involved by size expanded gene families

| Level 1 KEGG pathway | Level 2 KEGG pathway | Number of gene family | Detail of gene family (ID) |
| --- | --- | --- | --- |
| Cellular Processes | Cell growth and death | 2 | 6680;2948 |
| Cellular Processes | Transport and catabolism | 1 | 2967 |
| Environmental Information Processing | Signal transduction | 3 | 3683;2654;6680 |
| Environmental Information Processing | Signaling molecules and interaction | 2 | 2654;6680 |
| Genetic Information Processing | Folding, sorting and degradation | 1 | 794 |
| Metabolism | Global and overview maps | 2 | 9457;3404 |
| Metabolism | Glycan biosynthesis and metabolism | 2 | 9457;3404 |
| Organismal Systems | Development | 2 | 1171;6680 |
| Organismal Systems | Digestive system | 1 | 3933 |
| Organismal Systems | Endocrine system | 1 | 3683 |
| Organismal Systems | Environmental adaptation | 1 | 2654 |
| Organismal Systems | Immune system | 1 | 6680 |
| Organismal Systems | Nervous system | 1 | 2654 |
| Organismal Systems | Sensory system | 1 | 546 |

**Supplementary table 15.**
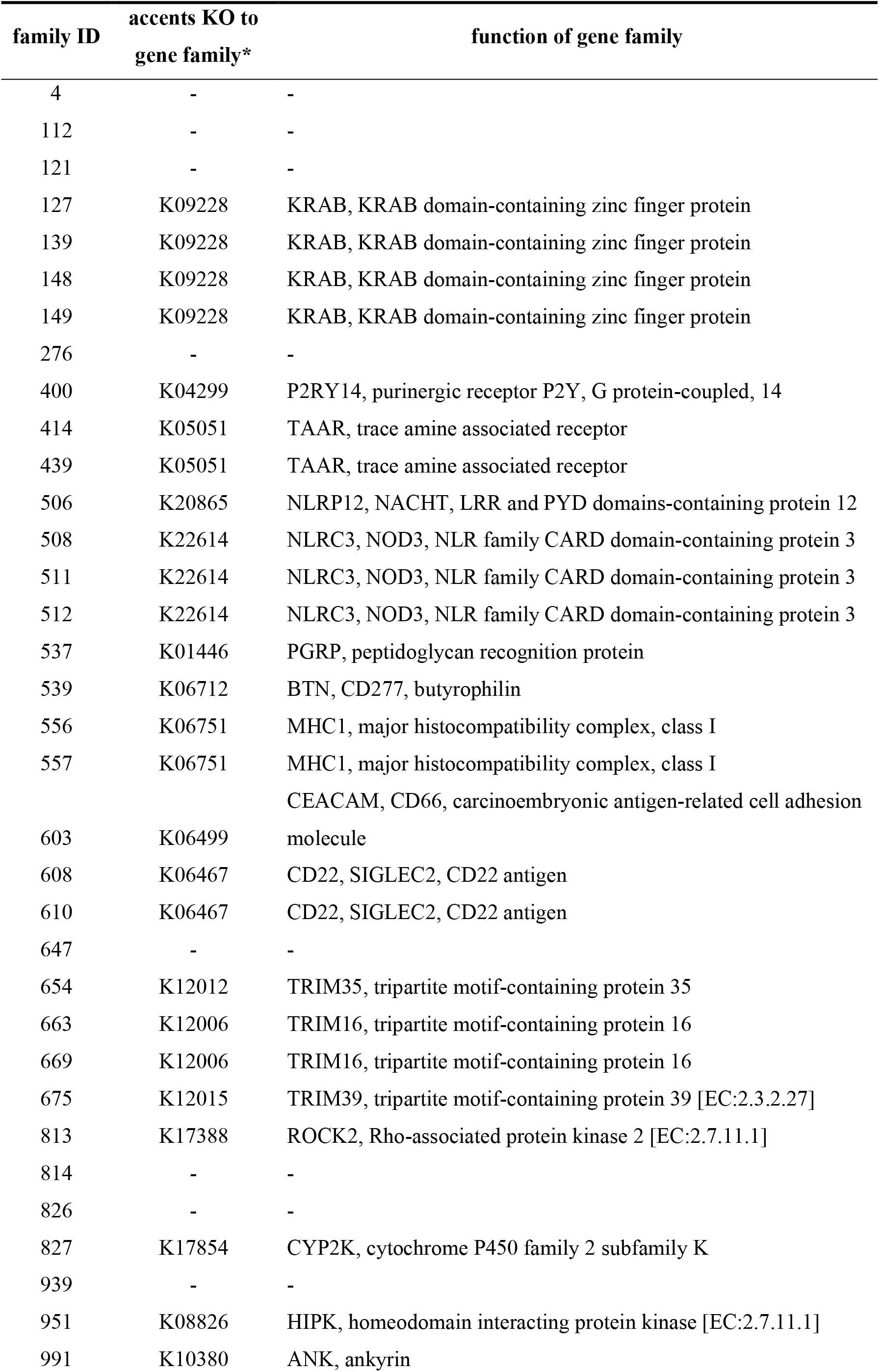
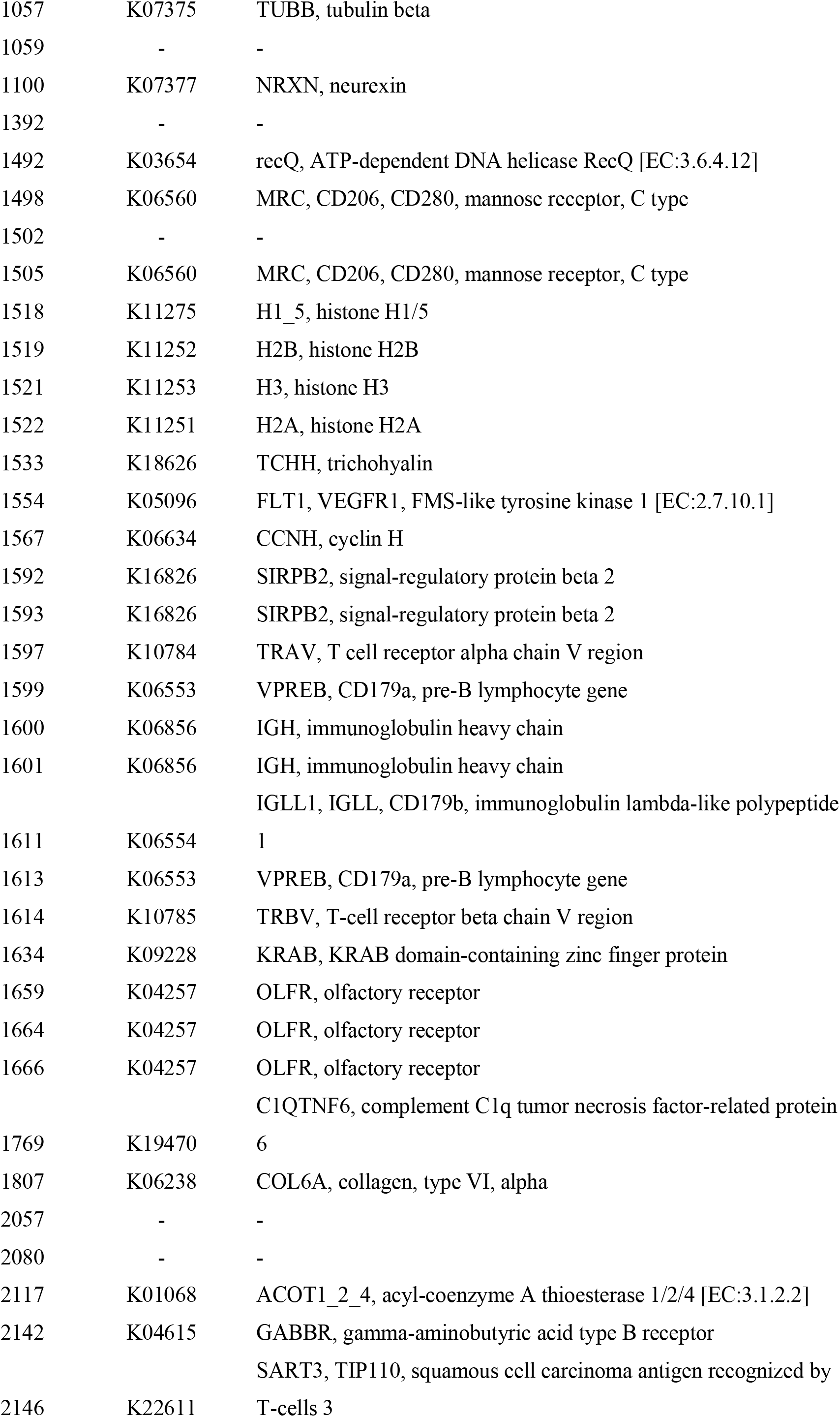
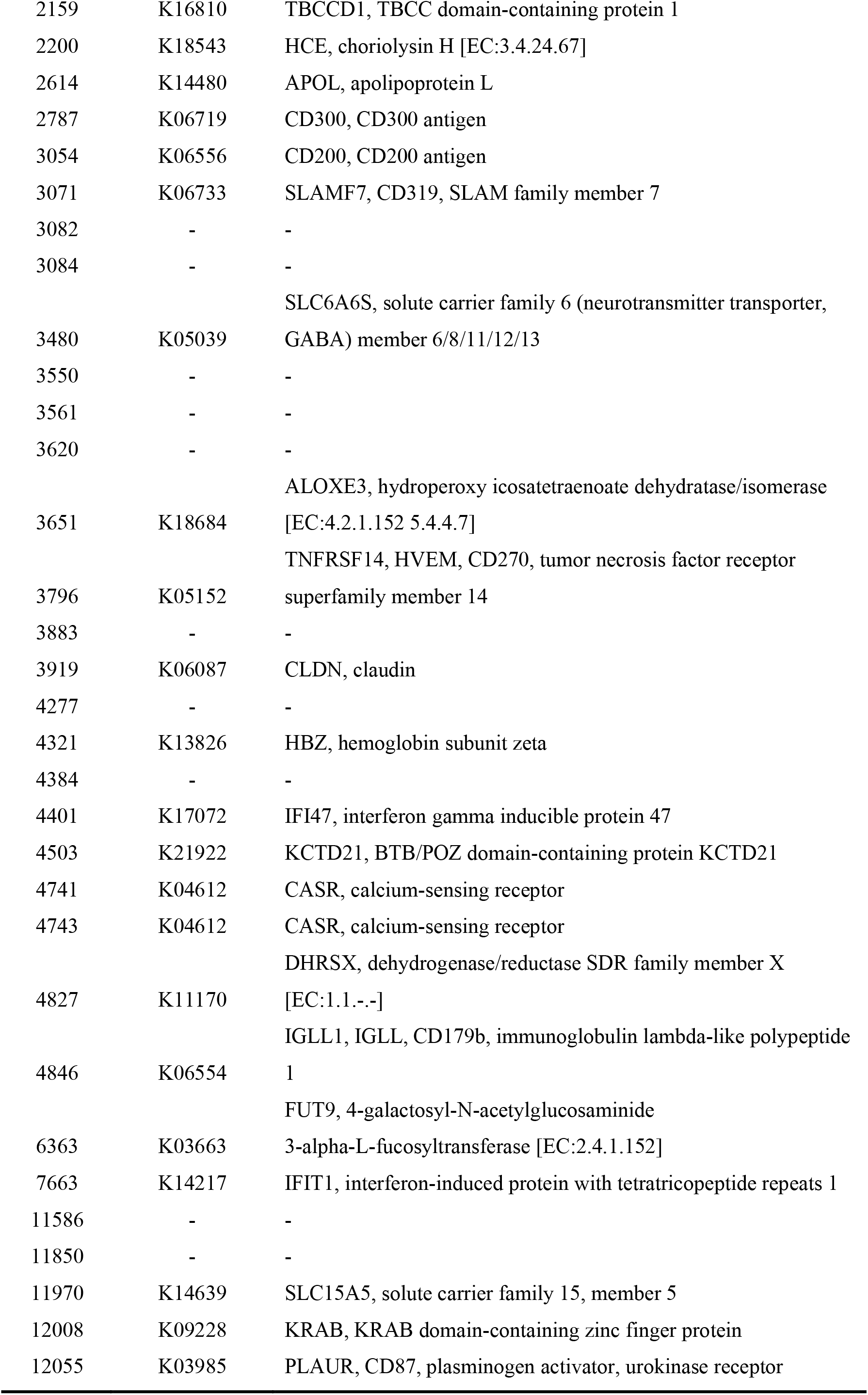
The detail for contracted gene families.

